# Development of a Novel SARS-CoV-2 Immune Complex Vaccine Candidate (CRCx) with Broad Immune Responses: A Preclinical Trial in Animal Model

**DOI:** 10.1101/2022.05.27.493693

**Authors:** Sherif Salah, Abdula Mubarki, Khalid Zayed, Khaled Omar

## Abstract

**Background:** The ongoing pandemic of COVID-19, caused by severe acute respiratory syndrome coronavirus 2 (SARS-CoV-2), poses a serious threat to global public health and imposes a severe burden on the entire human population. Faced with a virus that can mutate its structure while immunity is incapacitated, a need to develop a universal vaccine that can boost immunity to coronaviruses is highly needed.

**Design:** Five formulations of two types (CRCx2 and CRCx3) of immune complexes with an immunogen adjuvant were evaluated in a mouse model as candidate SARS CoV-2 vaccines in a pretrial prior to clinical trials in humans. CRCx3 comprises 3 different formulas and CRCx2 comprises 2. Balb/c mice were vaccinated intraperitoneally on days 0/7 with a high or low dose of CRCx2 or on days 0/7/14 with a high, medium, or low dose of CRCx3 series, and their blood was sampled for serum antibody measurements. Mice were challenged with live virus after immunization with either vaccine to evaluate prophylaxis ability or treated with them after challenge to evaluate therapeutic ability on day 15. Immunological markers and histopathological studies as well as titration of neutralizing antibodies to the vaccines were evaluated and analyzed.

**Results:** CRCx 3 and CRCx 2 vaccine candidates induced elevated levels of positive neutralizing antibodies as well as a cellular immune response with safety, efficient productivity, and good genetic stability for vaccine manufacturing to provide protection against SARS-CoV-2 with relatively higher levels with the high dose CRCx2 candidate combination.

**Conclusions:** Highly efficient protection and therapeutic effect against SARS-CoV-2 were obtained with a double-dose immunization schedule spaced at 7-day intervals using injections 0.25 of or 0.40 ml of CRCx2 vaccine formulations with a 25-mm needle. These results support further evaluation of CRCx in a clinical trial on humans.

## Introduction

Amid the severe acute respiratory syndrome coronavirus 2 COVID-19 (SARS-CoV-2) pandemic and its resultant morbidity and mortality, the need for safe and efficacious vaccines that induce protective and long-lived immune responses arises. Severe acute respiratory syndrome coronavirus (SARS-CoV) emerged in 2003, the Middle East Respiratory Syndrome Coronavirus (MERS-CoV) in late 2012, and COVID-19 (SARS-CoV-2), which is caused by a new positive-strand RNA coronavirus, in late 2019 after which it was declared a pandemic situation rapidly spreading worldwide on March 11, 2020, by the World Health Organization (WHO Director-General [1], [2], [3]. Globally, as of 7 April 2022, there have been 493,392,853 confirmed cases of COVID-19, including 6,165,833 deaths, and as of 5 April 2022, a total of 11,250,782,214 vaccine doses have been administered [4].

All three viral jumps belong to the *Coronaviridae* family whose classification recognizes 39 species in 27 subgenera [5], [6]. Of these, at least 7 coronaviruses are known to cause respiratory infections in humans of which four can cause common cold-like symptoms, and those that infect animals can evolve and become infectious to humans [6]. Coronaviruses’ genomes encode a number of major structural proteins such as spike (S), envelope (E), membrane (M), and nucleocapsid (N) proteins as well as approximately sixteen nonstructural proteins (nsp1–16) and five to eight accessory proteins [7]. Among them, the spike (S) protein plays an essential role in viral attachment, fusion, entry, and transmission and is the common target antigen for neutralizing antibodies and vaccine development [8]. The precise strategy used by coronaviruses for genome replication is not yet known, but many features have been established [9]. Several vaccine platforms and types of conventional vaccines are employed in vaccine development, each presenting several advantages and disadvantages in the race to obtain good results and safety [10], [11], [12], [13], [14], [15], [16], [17], [18], [19].

At present, there are more than 64 vaccine candidates, of which thirteen are being tested in Phase 3 clinical trials, most of them aiming to induce neutralizing antibodies against the spike protein (S) [10]. The relatively high speed of the development of vaccines makes it a promising approach to COVID-19 prevention; but, emerging evidence has shown antibody-dependent enhancement (ADE) in SARS-CoV infection [20], [21], suggesting that specific attention should be paid to safety evaluation in developing coronaviruses vaccines. In addition, it is still unclear whether the FDA-approved vaccines do block infection and virus shedding or only alleviate symptoms [22]. It has been reported that the probabilities of reinfection after the use of diverse vaccines have not been resolved or prevented [23]. Moreover, more reports were recorded of the possibility of serious side effects after the first and second doses [24], [25], [26], [27], [28], [29], [30], [31].

After SARS-CoV-2 infection, several categories of antibodies circulate in the blood including IgG, IgM, and IgA; mainly targeting the spike (S) protein and the nucleoprotein (NP). NP protein is abundant and highly expressed; however, due to its biological function, it seems to be unlikely that antibodies against NP have neutralizing activity [32]. On the other hand, the S protein contains the receptor-binding domain (RBD), which mediates the binding to the host cell through the human Angiotensin-Converting Enzyme 2 (ACE2) and the fusion of viral and cellular membranes [33], [34]. Generally, vaccines actively stimulate the immune system to produce antigen-specific neutralizing antibodies (NAbs) to prevent or treat disease. Understanding the structure and function of these proteins, particularly of the S protein and its RBD, provides a basis for the rational design and development of SARS-CoV-2-specific neutralizing antibodies (NAbs) for prophylactic prevention and treatment of COVID-19 [35]. Accordingly, most of the recent vaccines for COVID-19 that employ injection of viral antigens or viral gene sequences aim at inducing neutralizing antibodies against the viral spike protein (S), preventing uptake through the human ACE2 receptor; and thus, blocking infection [36], [37].

Studies of the immune mechanisms initiated in cells by Fc–FcR interactions presented the immune regulatory roles of antigen-antibody immune complex (IC) as a double-edged blade. Despite their potential to cause pathological effects, ICs have been explored as preventive as well as therapeutic vaccines, first in poultry breeding, and later in human diseases [38]. The first application of an IC vaccine was in the prevention of infectious bursa disease (IBD) in poultry and the first glimmer of success with an IC vaccine in HIV infections was reported by Hioe et al. (2009) [39]. When healthy adults were immunized with a seasonal trivalent influenza vaccine (TIV), IC showed that the abundance of sialylated Fc (sFc) on the anti-HA IgGs affected induction of the early plasmablast response and correlated with vaccine efficacy [40]. HBsAg-HBIG IC as a therapeutic vaccine for chronic HBV infection (CHB) was based on the concept that the Fc fragment of antibodies in the IC could interact with Fc receptors on DCs cells, and initiate more effective immune responses in hosts [41]. In mice, ICs were also used for Ebola [42] and the dengue virus [43]. Finally, a potential candidate IC for cancer patients is an IL-2-anti-IL-2 complex (IL-2 IC), that may extend IL-2 bioactivity from hours to days [44].

The rapid developments from studies on antibody structures and functions, genetic engineering technology for mass production of proteins, and novel methods of applying therapeutic antibodies have inspired more interest in neutralizing antibodies (NAbs). NAbs with high specificity, strong affinity to target proteins and low toxicity have been used to treat viral infections caused by Ebola virus, cytomegalovirus, influenza virus, human immunodeficiency virus (HIV), and respiratory syncytial virus [35]. NAbs generally exist in the body for a short time, and their treatment efficacy depends on many factors, including their titer, amount, specificity, and half-life [35]. Most approved vaccines against COVID-19 have traditionally focused on the induction of strong protective neutralizing antibodies against the target pathogen to confer long-term immunity in vaccinated individuals to protect the body from the risk of infection or recurrence [45]. These antibodies will prevent uptake through the human ACE-2 receptor, thus, hindering virus entry and, therefore, may comprise the potential key for future protection against COVID-19.

On the other hand, the degree to which SARS-CoV-2 will adapt to evade neutralizing antibodies is still unclear [46]. NAbs may carry a pathogenic effect that opposes its protective role as with ADE. Moreover, coronavirus may use these NAbs to mask a proportion of corresponding antigens in an antigen/neutralizing antibody immune-complex form (Ag/NAbs) for a long time, thus, preventing its attack by CD8+ cytotoxic T cells. The SARS-Cov-2 virus has unusual two different pathways that serve one another. First, targeting two types of cells in our bodies, the first and major target are the CD4+ T-helper cells, some of which the virus signal will cause to mutate into another form similar to CD8+ cells but with different physiological behavior. Meanwhile, the CD4+ cells will excite the B-cells to produce specific protective antibodies for every specific viral antigen. Second, the virus will invade its secondary target cells, which are the upper and lower respiratory tract cells, and starts replication. By the influence of the CD4+ cells on the secondary cells, the virus will be released in the form of separated viral particles that will be protected by the specific antibodies produced and create circulating immune complexes. Again, these viral antigens will get the chance to be released and infect other cells when the CD8+ cells are suppressed by the newly mutated CD8+ due to the dual immune response (See Supplement: Fig. 1). In addition, our humoral immune cells can produce positive or negative neutralizing antibodies and, in the case of Covid-19, the extrinsic immune complexes (IC_A_, IC_B_, and IC_C_) are completely different from the existing intrinsic circulating immune complexes that comprise coronavirus antigens; i.e. S, M, and N and their specific neutralizing antibodies IC_1_, IC_2_, and IC_3_. We believe that this antibody variation for the same antigen may be the reason behind the failure of many vaccines in inducing long-term immunity; however, it may as well be the cornerstone for producing highly effective vaccines.

**Figure 1:**
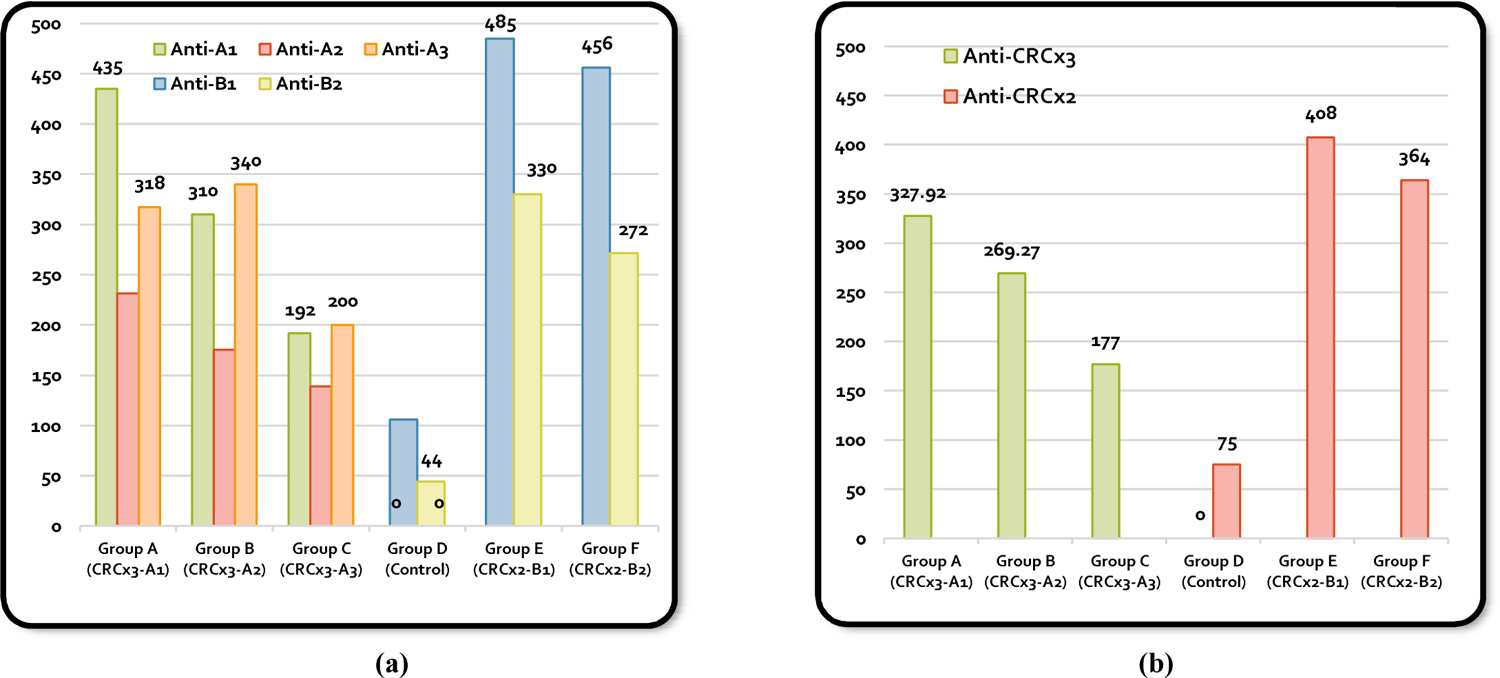
Mean neutralizing-antibody responses to the SARS-COV2 CRCx protective vaccines in vaccinated mice groups. (a) Individual neutralizing antibodies. (b) Total neutralizing antibodies.

First, we hypothesized that coupling antigen/nonspecific-antibody in one form, making it different from an existing intrinsic circulating immune complex (CIC) that shares antigen/specific-antibody may change the self-tolerance. This relates to the lack of immune response to the intrinsic CIC because of central (thymic selection) or peripheral (lack of co-stimulation) tolerance. We had wanted to whether if we inject an extrinsic noncomplex trigger that is relatively but not exactly similar to the intrinsic CIC, it can initiate a specific immune recognition of the tolerated CIC. Accordingly, for the first time in research for SARS-Cov-2 vaccines, we have undertaken this study to develop a new *in vitro* immune complex vaccine. Based on our novel postulation, our candidate vaccine (CRCx) should be able to provide both protective and therapeutic measures. In this report, we present the preclinical trial of two types or combinations of peptide immune complex SARS-CoV-2 vaccine candidates; collectively named CRCx vaccines. The patient will be injected with a combination, which contains either each specific viral antigen with its nonspecific antibodies (CRCx3) or with its specific antibodies (CRCx2).

They are designed to cause interactions that will trigger a series of immune-regulatory responses involving both the innate and adaptive immune systems, including cross-presentation of antigens, activation of CD8+ T-cells and CD4+ T-cells, phagocytosis, complement-mediated antibody-dependent cellular cytotoxicity (ADCC) and complement-dependent cytotoxicity (CDC). The cytotoxic cells use both mechanisms to destroy viruses. They will provoke CD8+ T-cells to secrete IFN-γ that will destroy the foreign complex as well as any other similar complex, thus, stopping the communication between CD4+ and upper and lower respiratory tract cells and interfering with the antigen production. Meanwhile, CD8+ cells will also block the virus’s tendency to infect other CD4+ cells and, accordingly, inhibit its mutation process. Finally, the immune system will be liberated from confusion created by the neutralizing antibodies and should be able to recognize and develop a positive immune response against the infection (See Supplement: Fig. 2). In our study, we demonstrate that this type of vaccine can achieve potency, safety, a long-lasting immune response, and boosted memory of coronaviruses. We wish that this is a new hope appearing on the horizon and that total victory against this virus is very near.

**Figure 2:**
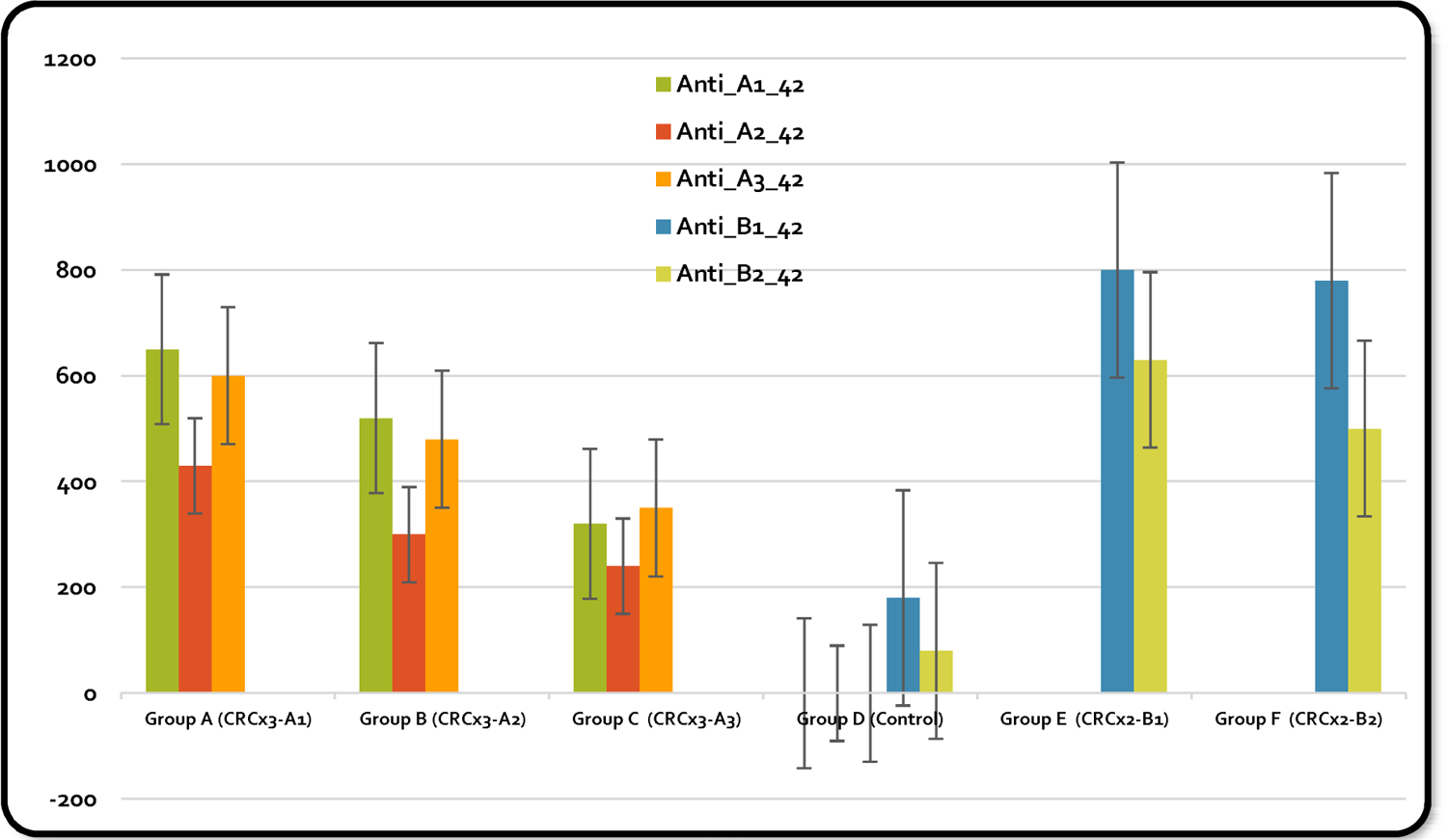
NAbs levels after day 42 when seroconversion was completed.

## 1. Materials and Methods

### 1.1. Facility and Ethics Statement

All the experiments with infectious live SARS-CoV-2 were performed in a biosafety level 3 (BSL3) and animal biosafety level 3 (ABSL3)-enhanced facility. This study was performed in accordance with the guidelines for the care and use of laboratory animals published and approved by the Committee for Ethics on Animal Experiments and the Committee for Animal Biosafety Level 3 Research of the Egyptian Military Scientific Commission.

### 1.2. Isolation of Viral Strain

Type VERO E6 cell line [from African Vervet monkey ‘*Cercopithecus aethiops*’ kidney cells (ATCC)], as a WHO-certified cell line for vaccine production and is routinely used for microorganism isolation or various diverse research projects in the facility, was used to replicate the viral strains. SARS-CoV-2 strains were screened to find an optimal viral seed after isolation from nasopharyngeal swabs of a positively-testing patient with viral pneumonia showing severe clinical demonstrations such as fever, seizure, muscle cramp, and reduction of arterial oxygen saturation as per Pavel *et al.* (2020) [47]. The rate of infection of the patient was evaluated via the qRT-PCR method. The lower the Ct (cycle threshold) of the qRT-PCR method, the higher the amount of viral load in the patient. One of the isolated strains from Vero cells that replicated the most and resulted in the highest virus yields among other strains (Ct < 20) was chosen for isolation of the viral strain.

### 1.3. Viral Titration

For efficient growth of viral stock in VERO cells, the isolated virus was first plaque purified and passaged once in VERO cells to generate the P1 stock. After this, three other passages were performed to generate the P2 to P4 stocks. Pre-cultured Vero cells were infected with an isolated viral strain in 96-well flat-bottom microplates (Dutscher). Then, a ten-step serial dilution of the purified viral replicates stock was prepared. Vero E6 cells with an average population of 10^4^ cells were cultured for 18–24 h in each well in the growth medium [Eagle’s Minimum Essential Medium (EMEM) (Lonza Bioscience) +L serum (FBS) at 37 °C+5% CO_2_], without antibiotics. A hundred microliter of each decuple (i.e., ten-fold) dilution steps was transferred to each well of the 96-well plates. The cells were observed daily for SARS-CoV-2 specific cytopathic effects (CPE) for 7 days using a Digital Inverted Fluorescence Microscope with a × 10 objective.

The supernatants were collected on day 0 and day 7 and viral titration was calculated using reverse transcription-polymerase chain reaction (RT-PCR) as per Amrane *et al*. [48]. The range for Delta Ct (ΔCt), the difference of Ct between day 0 and Day 7, for each cell type was calculated and done in triplicate. The cells for which a CPE effect was observed were incubated with 10-fold dilutions of different viruses and incubated for another 7 days at 37 °C. Each condition was performed in quadruplicate. The ΔCt (D_0_–D_7_ with Dil. 10^−1^) for VERO E6 cells ranged between 3.95-3.98, while for Dil. 10^−4^, it ranged between 13.5-16.2. The Median Tissue Culture Infectious Dose (TCID_50_) was determined by the Reed and Muench method [49] (See Supplement: Fig. 3). At multiplicities of infection (MOI) of 0.001-0.01, multiplication kinetics analysis of the P4 stock in VERO cells showed that the stock replicated efficiently and yielded a peak titer of 5 log^10^ TCID_50_/mL by 3 days after infection (dpi). The genomic content of Vero cells of each well with the minimal number of plaques was extracted for further molecular characterization. This process was repeated to complete four passages and a single plaque was obtained and its molecular identity was approved for viral challenge experiments.

**Figure 3:**
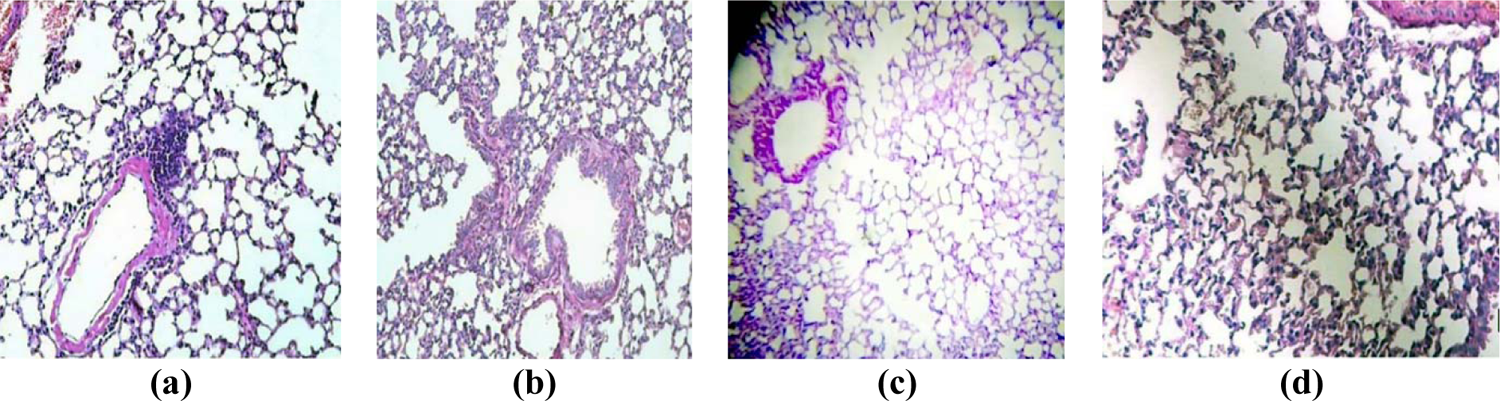
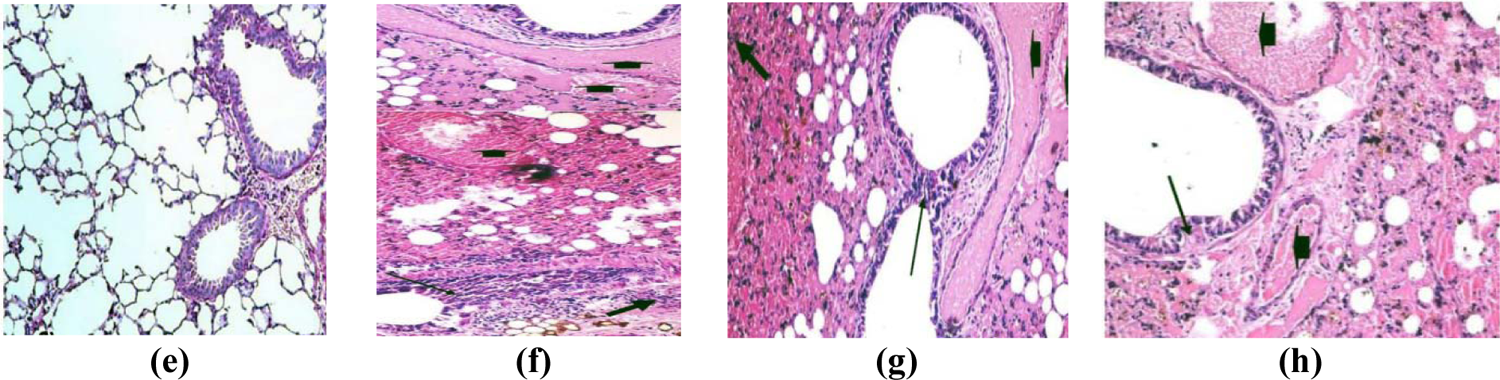
Normal lung tissue (a). Vaccinated mice with CRCx3 (A1, A2, A3) (b) CRCx2 (B1, B2) vaccinated mice (c) Test on 42 days (d) Post viral challenge showing normal lung tissues in CRCx3 vaccinated mice (e) Post viral challenge showing normal lung tissues in CRCx2 vaccinated mice. (f), (g) & (h) Unvaccinated mice lung at 14 & 28 days post-challenge test showing diffuse thickening in the interstitial tissue, perialveolar blood capillaries congestion, and lymphocytic cells infiltrations.

### 1.4. Preparation of Antigen

Different SARS-CoV-2 coronavirus antigen concentrations (5, 10, 20, 40, and 80 μg/ml) of spike protein (S1) (by ProSci Inc.), nucleocapsid protein, and membrane protein peptide (by AMSBIO) were procured, diluted with 1 ml of phosphate-buffered saline (PBS); i.e. (1/400, 1/200, 1/100, 1/50, 1/25) and incubated at room temperature for at least 2 hours.

### 1.5. Preparation of Antibodies

Different SARS-CoV-2 antibodies concentrations (5, 10, 20, 40, and 80 μg/ml) antibody (by ProSci Inc.), nucleocapsid antibody, and membrane antibody (by AMSBIO) were procured, diluted with 1 ml of PBS; i.e. (1/400, 1/200, 1/100, 1/50, 1/25) and incubated at room temperature for at least 2 hours.

### 1.6. Preparation of Immune Complexes

Two different modes of immune complexes formulation were used in which mixing of a specific concentration of an antigen with its relevant concentration of an antibody took place and named A1, A2, A3, B1, and B2, where A1, A2, and A3 constitute the CRCx3 series and B1 and B2 constitute the CRCx2 series. The first mode (a non-specific mixture composed of coronavirus antigens and their non-specific antibodies) included SARS-CoV-2 spike protein (S1 subunit) with anti-nucleocapsid antibodies (A1), nucleocapsid antigen (N) with anti-membrane antibodies (A2) and membrane antigen (M) with anti-spike antibodies (A3). The resultant three complexes were collectively dubbed CRCx-3. The second mode (a specific mix composed of coronavirus antigens and their specific antibodies) included spike antigen (S1) with anti-spike antibodies (B1), and nucleocapsid antigen (N) with anti-nucleocapsid antibodies (B2). The resultant two complexes were collectively dubbed CRCx-2.

The different preparations of antigens and antibodies were mixed, allowed to react, and incubated for 6 hours at room temperature. Then, preparations were centrifuged at 2500 rpm for 10 minutes and the supernatant was discarded. For identification studies of the anti-complex antibodies, a fraction of the obtained sediment of every preparation was sampled. Identification and assessment of non-purified and purified vaccines were carried out by sodium dodecyl sulfate-polyacrylamide gel electrophoresis (SDS-PAGE) and Western blotting (WB) analysis. To finalize the complex preparations (A1, A2, A3, B1, and B2) for inducing anti-complex antibodies production in immunized rabbits, 3 ml of the adjuvant goat IgG anti-human IgM Fc (5 ng/ml) (by ABCAM, Cambridge, UK) was dissolved in human albumin and phosphate-buffered normal saline added to the remainder of the sediments and packaged in 1 ml vials of injectable solution with labels (See Supplement: Fig. 4).

**Figure 4:**
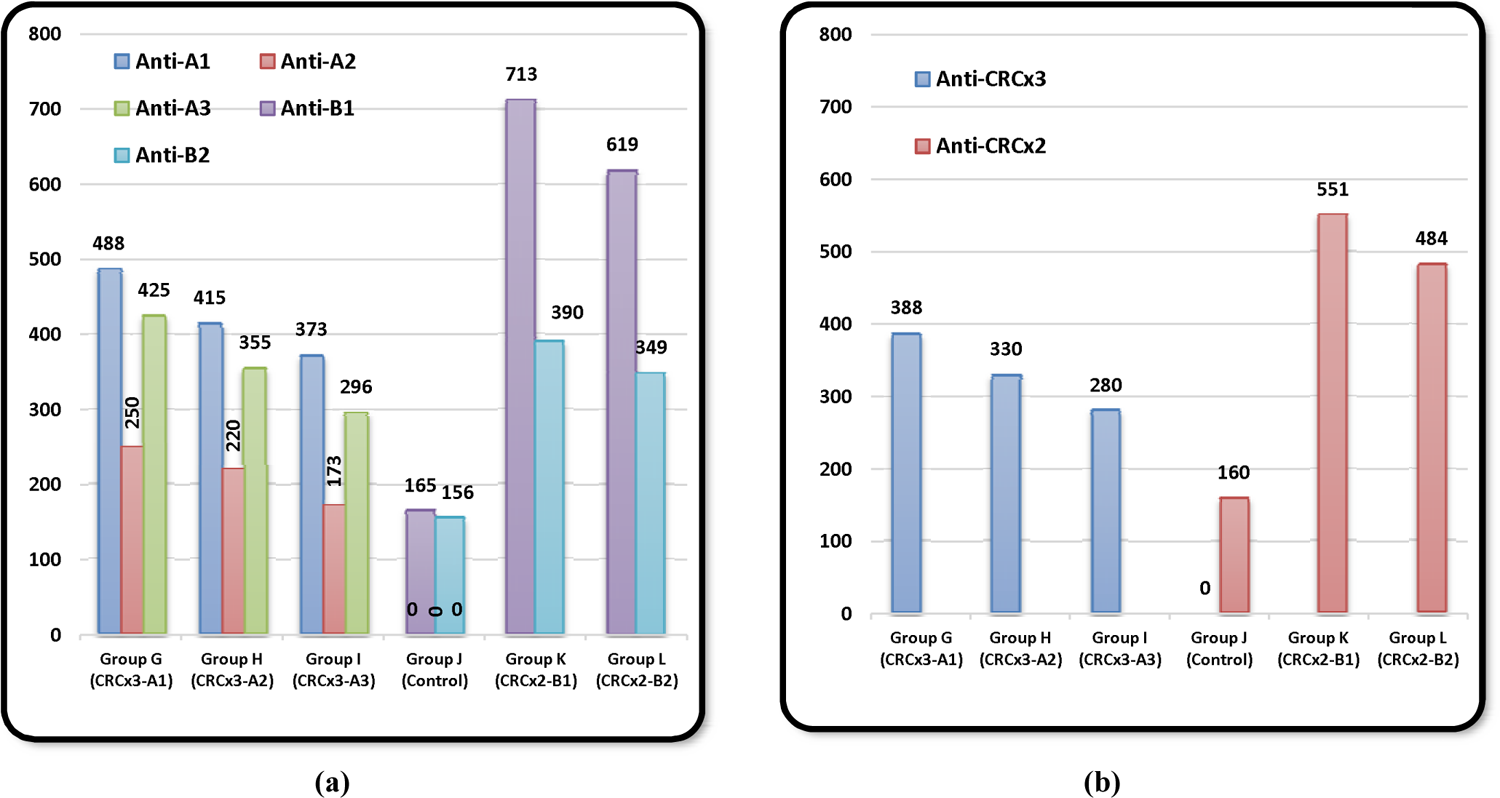
Mean neutralizing-antibody responses to the SARS-COV2 CRCx therapeutic vaccines in vaccinated mice groups. (a) Individual neutralizing antibodies. (b) Total vaccine groups: CRCx3 & CRCx2

### 1.7. Animal Studies

In this study, groups of BALB/c transgenic mice, rats, rabbits, and guinea pigs were recruited. In all experiments, all animal care was conducted under the guidelines for animal experiments and performed as specified in regulations, which describe animal protection and working with laboratory animals in the country. Throughout the experiments in this study, strict quality control and quality assurance measures were followed according to the institution and laboratory regulations. All efforts were made to minimize animal suffering. All animals were observed for morbidity and mortality, overt signs of toxicity (including abstinence from food and water), and any signs of distress throughout the studies (See Supplement: Table 1).

**Table 1:** Paired t-test results of correlation and differences between vaccinated groups (A, B & C) and control group (D) as regards NAbs levels.

| Paired Samples Descriptive Statistics |  |  |  |  | Correlations |  | Differences |  |  |  |  |  |  |
| --- | --- | --- | --- | --- | --- | --- | --- | --- | --- | --- | --- | --- | --- |
| Groups | Mean | N | Std. Dev. | Std. Error Mean | Cor. | Sig.* | Mean | Std. Dev. | Std. Error Mean | Lower 95% CI Dif | Upper 95% CI Dif | t | Sig. (2-tailed)* |
| Group A | 327.9 | 16 | 179.6 | 44.9 |  |  | 327.9 | 179.6 | 327.9 | 179.6 | 44.9 | 7.301 | 0.000 |
| Group D | 0.0 | 16 | 0.0 | 0.0 |  |  |  |  |  |  |  |  |  |
| Group B | 270.5 | 15 | 127.1 | 32.8 |  |  | 270.5 | 127.1 | 270.5 | 127.1 | 32.8 | 8.242 | 0.000 |
| Group D | 0.0 | 15 | 0.0 | 0.0 |  |  |  |  |  |  |  |  |  |
| Group C | 177.0 | 16 | 96.0 | 24.0 |  |  | 177.0 | 96.0 | 177.0 | 96.0 | 24.0 | 7.372 | 0.000 |
| Group D | 0.0 | 16 | 0.0 | 0.0 |  |  |  |  |  |  |  |  |  |
| Group E | 345.0 | 13 | 262.4 | 72.8 | 0.800 | 0.001 | 281.1 | 205.9 | 57.1 | 156.7 | 405.6 | 4.923 | 0.000 |
| Group D | 63.8 | 13 | 77.1 | 21.4 |  |  |  |  |  |  |  |  |  |
| Group F | 308.1 | 13 | 244.7 | 67.9 | 0.827 | 0.000 | 244.2 | 186.0 | 51.6 | 131.8 | 356.6 | 4.733 | 0.000 |
| Group D | 63.8 | 13 | 77.1 | 21.4 |  |  |  |  |  |  |  |  |  |

\*P-value is significant at 0.05 (2-tailed)

For the production of different IC antibodies, twenty-four male rabbits were divided into six groups (numbered 1 through 6), each comprising four rabbits. Groups 1 through 5 were injected with three intramuscular (0.25 ml) injections of one of the immune complexes: A1, A2, A3, B1, and B2, respectively for 25 days. Group number 6 acted as a control group and received only PBS. Six weeks later, all vaccinated animals were bled and, from their sera, anti-complex precipitated globulin antibodies were collected.

For prophylactic and therapeutic effects and immunogenicity studies of candidate vaccines, sixty BALB/c mice of six to eight weeks of age and mixed gender were obtained from Military Central Laboratories Animal Center where they were bred and maintained in a specific pathogen-free environment. All selected mice for the study were in good health and were not involved in any other experiment. Animals were housed in cages covered with barrier filters in a biosafety level 3 (ABSL3) enhanced facility with a 12-h light/dark cycle and access to food and water *ad libitum* in addition to environmental enrichment. The mice were maintained at an 18-28°C temperature and relative humidity of 55±15%. All mice were monitored daily to ensure that they were eating, drinking, and behaving normally as well as for care and health.

Mice were randomly divided into twelve groups, five mice per group, and classified into two categories. To obtain maximum antibody production from minimum antigen content in CRCx formulations, two approaches of injection were employed. In the first approach, which was designed to study the safety, efficacy, and efficient safe dose of the vaccines as protective or preventive vaccines, the first six groups of Balb/c mice tagged (A, B, C, D, E, and F), where group D, acting as a control group for this category, were given different regimens of intraperitoneal double/triple infusion of three different concentrations of high (HD): 0.4 ml/dose, medium (MD): 0.35 ml/dose, and low (LD): 0.25 ml/dose as explained below in trail 1. In the second approach, which was designed to study the safety, efficacy, and the efficient safe dose of the vaccines as therapeutic vaccines, the later six groups of Balb/c mice tagged (G, H, I, J, K, and L), where group J acting as a control group for this category, were given different regimens of intraperitoneal double/triple infusion of three different concentrations of high (HD): 0.4 ml/dose, medium (MD): 0.35 ml/dose, and low (LD): 0.25 ml/dose as explained below in trail 4.

In addition, for safety and toxicity studies of the vaccines, animal groups of rats, rabbits, and guinea pigs were used. Twenty-four mixed-gender rats were divided into four groups, three experimental (n=6 each) and a control group (n=6) for toxicity studies. Ten rabbits were divided into 5 groups (3 with n=3 and 2 with n=2) for vaccine pyrogenicity and local irritation studies. In addition, twenty-four male guinea pigs were divided into 4 groups (n=6 each) for active systemic anaphylaxis, repeated dose toxicity, and local irritation.

### 1.8. Post-immunization Production of Different IC Antibodies in Rabbits’ Sera

Precipitation of anti-complex antibodies was carried out by ammonium sulfate at half-saturation by adding 5 gm of solid ammonium sulfate to 15 ml of serum. Each group of rabbits produced 15 ml of immune serum (200 mg) containing antibodies to the immune complexes A1, A2, A3, B1, or B2.

The extracted insoluble antibodies are then separated by filtration and the solid material is dialyzed against 0.85% sodium chloride until free from the sulfate. A small amount of insoluble protein may appear during the dialysis, which can be removed by filtration with Whatman® qualitative filter paper, Grade 3. Then, the clear filtrate is added to the volume of the original serum sample. The supernatant fluids which contain the complex structures and the non-bound antibodies to these complexes are then discarded. The high molecular weights of antigen– antibody complexes help precipitation with ultracentrifugation. The amounts of anti-complex antibodies precipitated from each rabbit’s sample are then estimated according to the different concentrations of injected complexes mixtures by ELISA.

### 1.9. ELISA Assay of Produced Immune Complexes Antibodies

For quantitative titration of anti-immune-complexes produced from experimental rabbits’ sera to the different immune complexes formulae, an ELISA assay was performed by coating each one of five 96-well plates with a specific immune complex of A1, A2, A3, B1, or B2. Each ELISA Assay Kit (numbered 1 through 5) contains the key components (numbered A through F) required for the quantitative analysis of SARS-CoV-2 antigen-antibodies complexes concentrations in the sera of tested animals within the range of 5 to 200 μ /ml in a sandwich ELISA format. Monoclonal antibodies were selected as capture antibodies for each of the five immune complexes. The components in each kit were designed to be sufficient for the assay in 96-well ELISA plates.

Kits components included a component A, which is a coated plate (manufactured by Thermo Scientific, USA) with 50 μl/well of 5 μg/ml of the five different SARS-CoV-2 antigen-antibody complexes (A1, A2, A3, B1, or B2). Component B is an anti-complex-antibodies standard calibrator: 5, 10, 20, 30, 40, and 50 μg/ml (previously prepared from rabbits’ sera) for each of the immune complexes (A1, A2, A3, B1, or B2). Components C through F are the same for all 5 kits (Sigma-Aldrich) and are as follows: Component C is Horseradish Peroxidase (HRP)-Conjugated Anti-Rabbit IgG 25 μl, component D is a TMB Solution A (3, 3’, 5, 5’-tetra-methyl-benzidine) 15 ml, component E is a TMB Solution B (H_2_O_2_) 15 ml and component F is a TMB Stop Solution 30 ml.

### 1.10. Assessment of Immunogenicity and Efficient Doses of (CRCx2) and (CRCx3) as Prophylactic Vaccine Candidates

#### Trail 1

The BALB/c mice in the groups A, B, and C were intraperitoneally injected with various doses (0.25, 0.35, 0.40 ml /dose) of CRCx3 vaccines on days 0, 7, and 14 [primary (A1), booster (A2) and 2^nd^ booster (A3)]. Mice in group A were injected with a high dose (0.40 ml), and group B with a medium dose (0.35 ml) while mice in group C were injected with a low dose (0.25 ml). In addition, the BALB/c mice in groups E, and F were intraperitoneally injected with CRCx2 on days 0 and 7 [primary (B1) and booster (B2)], a high dose of (0.40 ml) for group E and a low dose (0.25 ml) for group F. Group D mice acted as controls and were injected with normal saline as placebo (See Supplement: Fig. 5).

**Figure 5:**
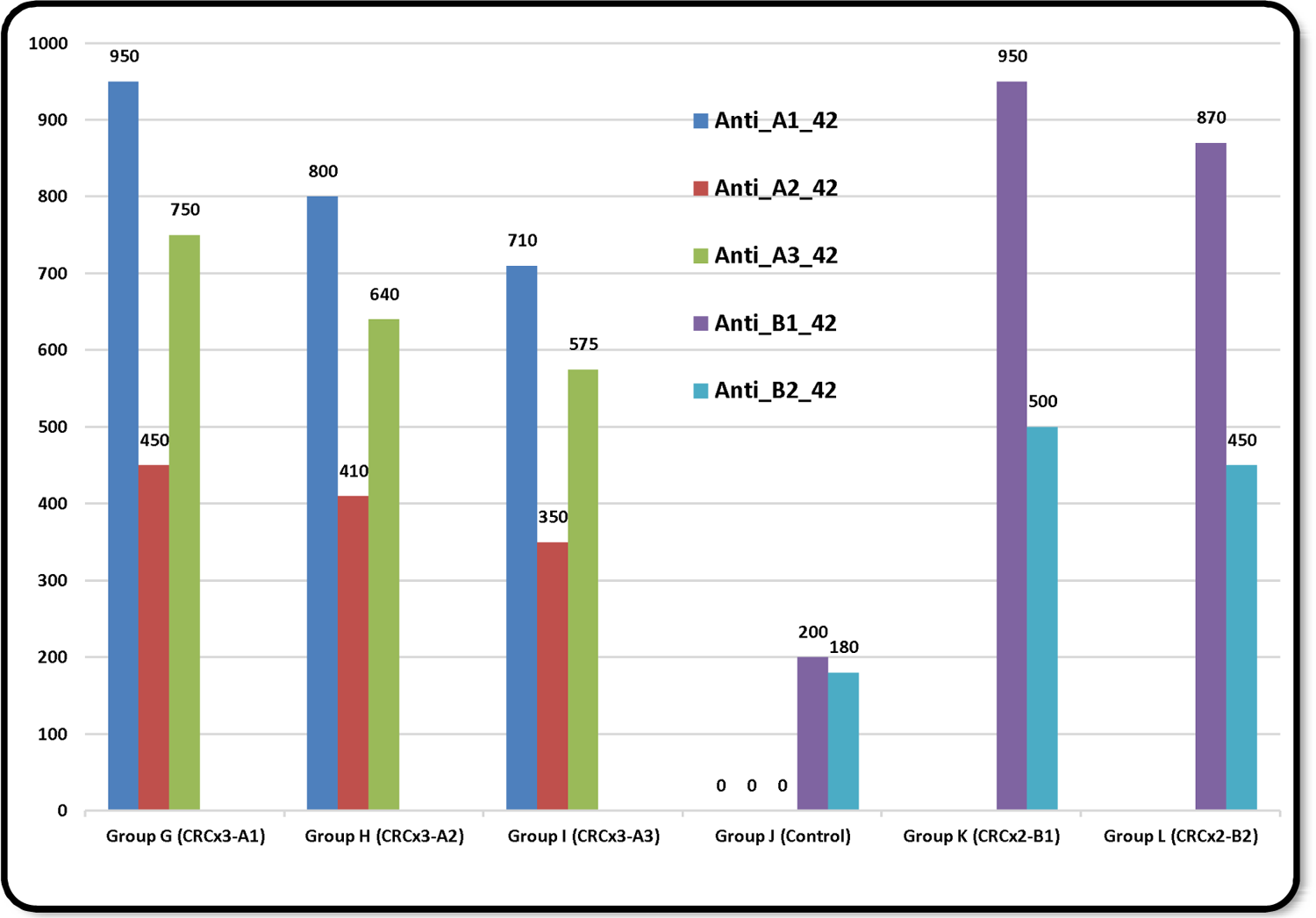
NAbs levels after day 42 when seroconversion completed.

#### Trail 2

For 14 days, starting on the day 15 from primary immunization in trail 1, the BALB/c mice of groups A, B, C, D, E, and F were intramuscularly challenged in the upper leg with a daily dose/mouse of 0.25 ml (total of 10^8^ IU) of live SARS-Cov-2 virus to which 5 IU of DNA polymerase was added. The virus used was taken from the working stock prepared in Vero E6 cultures as described above.

Blood samples were collected from each animal by retro-orbital (RO) route before immunization, and the serum was isolated the next day as a control. Also, from day zero, blood samples were taken on days 14, 28, 42 & 56 to measure serum immunological markers and anti-complexes antibodies. Sera optical density at 450 nm was measured and the levels of anti-CRCx3 and anti-CRCx2 neutralizing antibodies (NAbs) were evaluated. Clinical manifestation, body weight, and body temperature of the animals were monitored during and after immunization. In addition, Virus Neutralizing Test (VNT) in mice groups was performed. All mice were sacrificed on day 29 post-inoculation (dpi) and the lungs were used for viral load and pathological analysis.

### 1.11. Assessment of Therapeutic Efficacy of (CRCx2) and (CRCx3) as Vaccine Candidates

#### Trail 3

For 14 days, the BALB/c mice of groups G, H, I, J, K, and L were intramuscularly challenged in the upper leg with a daily dose/mouse of 0.25 ml (total of 10^8^ IU) of live SARS-Cov-2 virus to which 5 IU of DNA polymerase was added. The virus used was taken from the working stock prepared in VERO E6 cultures as described above.

#### Trail 4

The BALB/c mice in the groups G, H, and I were intraperitoneally injected with various doses (0.25, 0.35, 0.40 ml /dose) of CRCx3 vaccines on days 15, 22, and 29 [primary (A1), booster (A2), and 2^nd^ booster (A3)]. Mice in group G were injected with a high dose (0.40 ml), group H with a medium dose (0.35 ml) while mice in group I were injected with a low dose (0.25 ml). In addition, the BALB/c mice in groups K and L were intraperitoneally injected with CRCx2 on days 15 and 22 [primary (B1) and booster (B2)], a high dose of (0.40 ml) for group K, and a low dose (0.25 ml) for group L. Group J mice acted as controls and were injected with normal saline as placebo (See Supplement: Fig. 6).

**Figure 6:**
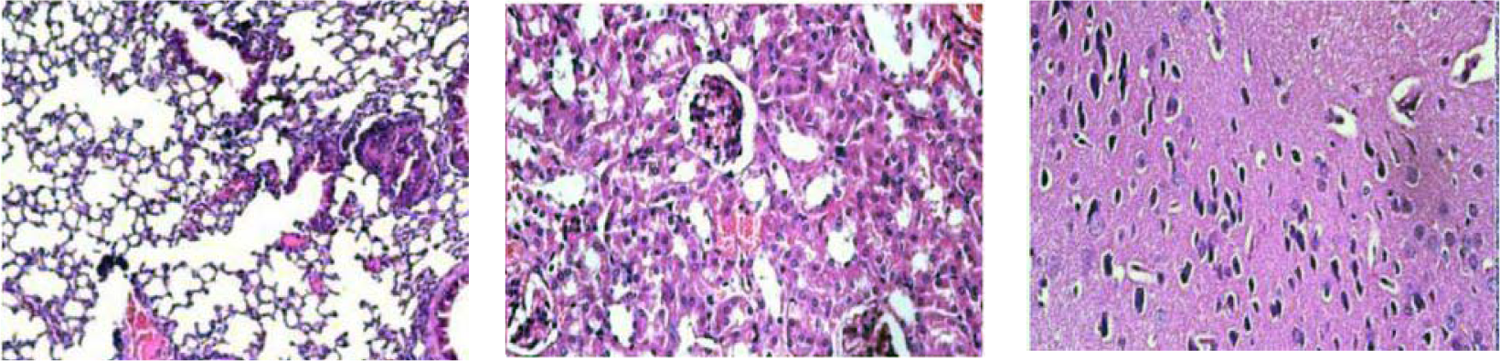
Normal organs’ tissues of vaccinated mice with two doses (Triple High=1.2 ml, Triple Medium=0.8 ml, or Triple Low=0.6 ml) of candidate inactivated SARS-COV2 vaccine CRCx (H&E X 100).

Blood samples were collected from each animal before inoculation. Also, from day zero, blood samples were taken on days 14, 28, 42 & 56 to measure serum immunological markers and anti-complexes antibodies. Sera optical density at 450 nm was measured and the levels of anti-CRCx3 and anti-CRCx2 neutralizing antibodies (NAbs) were evaluated. In addition, throat and anal swabs were obtained from mice on days 25 and 40 to evaluate the viral loads in tested groups. Clinical manifestation, body weight, and body temperature of the animals were monitored during and after inoculation. In addition, Virus Neutralizing Test (VNT) in mice groups was performed. All mice were sacrificed on day 29 post-immunization and the lungs were used for viral load and pathological analysis.

### 1.12. Biochemical and Immunological Assay

Serum samples were collected from all animals in the above-mentioned trails to evaluate the biochemical parameters and immunological markers. These included lymphocyte subset percentages (CD4+, CD8+ and CD4+/CD8+ ratio), quantitative inflammatory array (IL-1 alpha, IL-1 beta, IL-4, IL-6, IL-8, IL-10, IL-13, IFN-gamma, TNF alpha), D-dimer, C-Reactive Protein (CRP), Human Plasminogen Activator Inhibitor 1 (PAI1), LDH assay kit/cardiac troponin I enzyme, lactate dehydrogenase, serological detection of anti-CRCx2 and anti-CRCx3 complexes IgG and IgM, serum SARS-CoV-2 IgM and IgG, and nasotracheal or anal swabs for detection of post-inoculation qualitative SARS-CoV-2 RNA.

### 1.13. Evaluation of Safety of (CRCx2) and (CRCx3) as Vaccine Candidates

For determining the safety and/or single/repeated doses toxicity (including local irritation) of CRCx3 and CRCx2 as vaccine candidates, an experiment of double intramuscular injection in rats was administered to evaluate the possibility of vaccines’ acute toxicity. In this experiment, twenty-four rats were divided into four groups, three experimental (n=6 each, 3 of each gender) and one control group (n=6, 3 of each gender). The experimental groups were intramuscularly injected with two doses (Triple High=1.2 ml, Triple Medium=0.8 ml, or Triple Low=0.6 ml) of candidate vaccines, while the control group was infused with physiological saline, on days 0 and 7. All rats were monitored for 14 days to assure that there are no cases of death, any significant difference in weight or feeding state, and histopathologic changes after euthanasia (15 days after vaccination). The site of injection was also examined for erythema and edema at 24, 48, and 72 hours to check for local tolerance. All animals were examined for morbidity and mortality during the experimental period and were euthanized, necropsied, and the organs were dissected for macroscopic and microscopic evaluation.

In addition, for pyrogenicity assay, thirteen rabbits of approximate same weight were selected and divided into 5 groups (n=3 rabbit * 3 groups for CRCx3 and n=2 rabbit * 2 groups for CRCx2). Rabbits were deprived of food and only had access to water 24 hours before the injection. After one hour, the initial temperature of the animal was taken with a special rectal thermometer placed in the animal’s rectum. Normal doses (0.40, 0.35, and 0.25 ml) of each of the CRCx3 formulas were injected into the marginal vein of each rabbit’s ear in the first 3 groups, and doses (0.40 and 0.25 ml) of each CRCx2 formula in the remaining two groups. The post-injection body temperature of rabbits was measured in the first, second, and third hours post-injection.

Furthermore, acute systemic anaphylaxis due to CRCx was evaluated by intramuscular and intravenous injections in guinea pigs. Twenty-four male guinea pigs were divided into 4 groups (6 per group) as follows: a negative control group (physiological saline), a positive control group (human blood albumin, 20 µg/sensitization, 40 µg/stimulation), a low-dose group (0.1 x dose/sensitization, 0.2 x dose/stimulation), and a high-dose group (1 x dose/sensitization, 2 x dose/stimulation). Sensitization was performed on days 1, 3, and 5. The primary stimulation (intravenous excitation via foot) was performed on 3 out of 6 guinea pigs from each group on day 19, and secondary stimulation of the remaining animals of each group was performed on day 26.

### 1.14. Histopathology and Immunohistochemistry

Animal necropsies were performed according to a standard protocol. Tissues were fixed in formalin and embedded in paraffin blocks. 5 µm sections were stained with hematoxylin and eosin (H&E) and stored in 10% neutral-buffered formalin for 7 days before light microscopy examination for histopathological interpretation. 3 µm sections of lung paraffin blocks were used for immune-histochemical staining. The slides were stained with DAPI (5 µg/mL) after washing with PBS. Anti-CD4+ and Anti-CD8+ were the markers that were applied for the detection of immune cells in different parts of lung sections. Throughout each slide, 10 randomly chosen fields were evaluated in a high power field (HPF) for counting CD4+ and CD8+ lymphocytes by light microscopy (Nikon Eclipse E600, Japan).

### 1.15. Statistical Analysis

The SPSS software Version 25 was used to analyze the level of significance, using one-way ANOVA, paired t-test (2-tailed), and Pearson’s correlation methods. The confidence interval was set at 95% and *P* ≤ 0.05 was considered statistically significant. Neutralizing antibody titers, lung virus levels, and histopathologic changes scoring were averaged for each group of mice. Comparisons were conducted using parametric and nonparametric statistics as indicated.

### 1.2. Results

#### 2.2.1. Immunogenicity and Protection in Mice

First, to assess the humoral immunogenicity and function of CRCx as a prophylactic candidate, BALB/c mice were injected with triple combinations [0 (A1)/7 (A2)/14 (A3) days in groups A (High dose), B (Medium dose) & C (Low dose)] and double [0 (B1)/7 (B2) days in groups E (High dose) & F (Low dose)] immunization programs. In other words, the BALB/c mice in groups A & E received high (HD), B received medium (MD) and C & F received low (LD) doses of the candidate vaccines. Group D received normal saline as a placebo. The level of neutralization antibody (NAb) was evaluated after injection in each group. In detail, upon challenge by direct inoculation of 10^6^ TCID_50_ of SARS-CoV-2 into the vaccinated and control mice intramuscularly on day 15 (2 weeks after the primary, one week after 1^st^ booster or 1 day after 2^nd^ booster immunization dose). Clinically, CRCx3 mice in group A (4 of 5), in group B (2 of 5), and in group C (all 5) demonstrated elevations in temperature and became lethargic. Furthermore, in the CRCx2 mice group E and group F, all five mice demonstrated an elevation in temperature but soon became more active than normal with no signs of illness nor changes in body weights. In contrast, in the mock group (D), two mice died on day 25 post-immunization and the remaining three mice showed marked elevation in temperature, lethargy, weight loss, and sneezing. They also developed ruffled fur, a hunched posture, and rapid breathing.

In all immunized groups, all five mice showed a marked decrease in viral load by day 42 and became almost undetectable by day 56. However, after the virus challenge, the amount of viral load was undetectable in CRCx3 compared to the CRCx2 groups while the mock group showed a significantly high viral load. All the biochemical and immunological markers dramatically decreased by day 56. In general, even though by the end of day 14 (DPI), all animal group mice showed signs of pneumonia with marked elevation in body temperature, nervous manifestations, and marked elevations in all biochemical and biological markers, yet by 4 weeks’ time post-immunization, all vaccinated mice were highly protected against SARS-CoV-2 infection. There were no obvious weight loss changes among the vaccinated groups, inflammation, or any other observed adverse effects (See Supplement: Table 2, Fig. 7).

**Figure 7:**
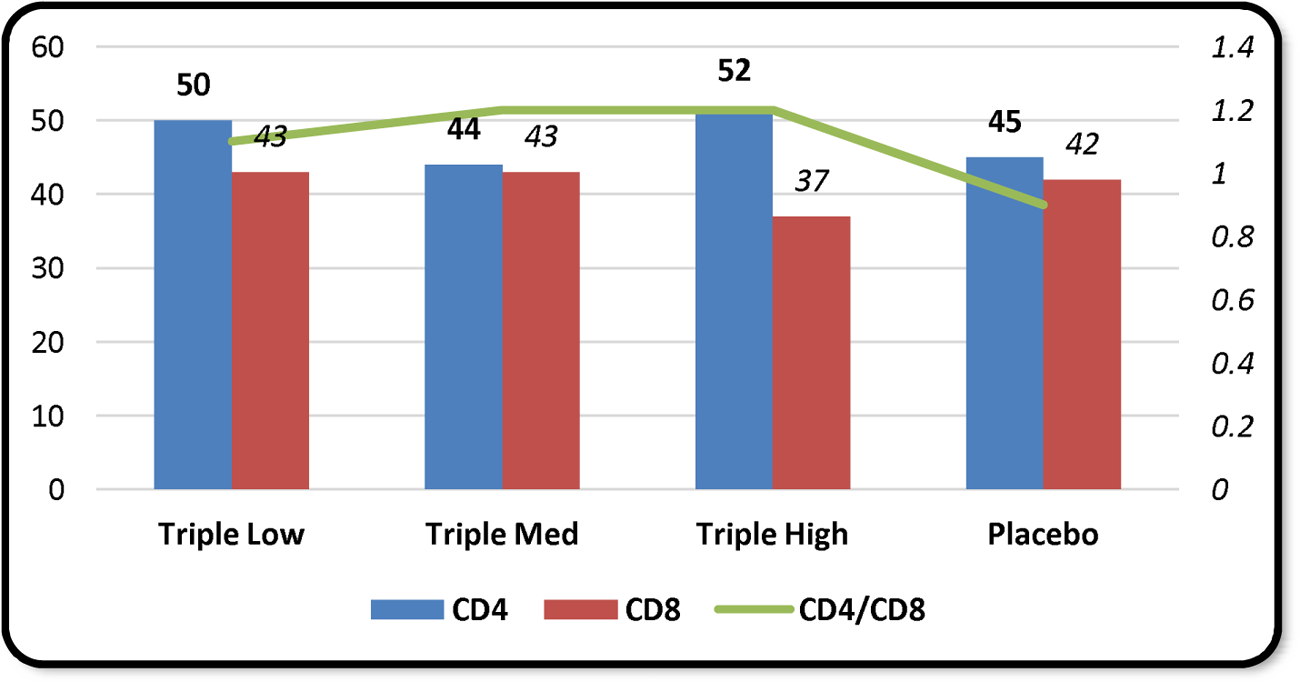
Cellular immunity evaluation after administration of a triple-low, -medium, and -high dosages of CRCx candidate vaccines in rats.

**Table 2:**
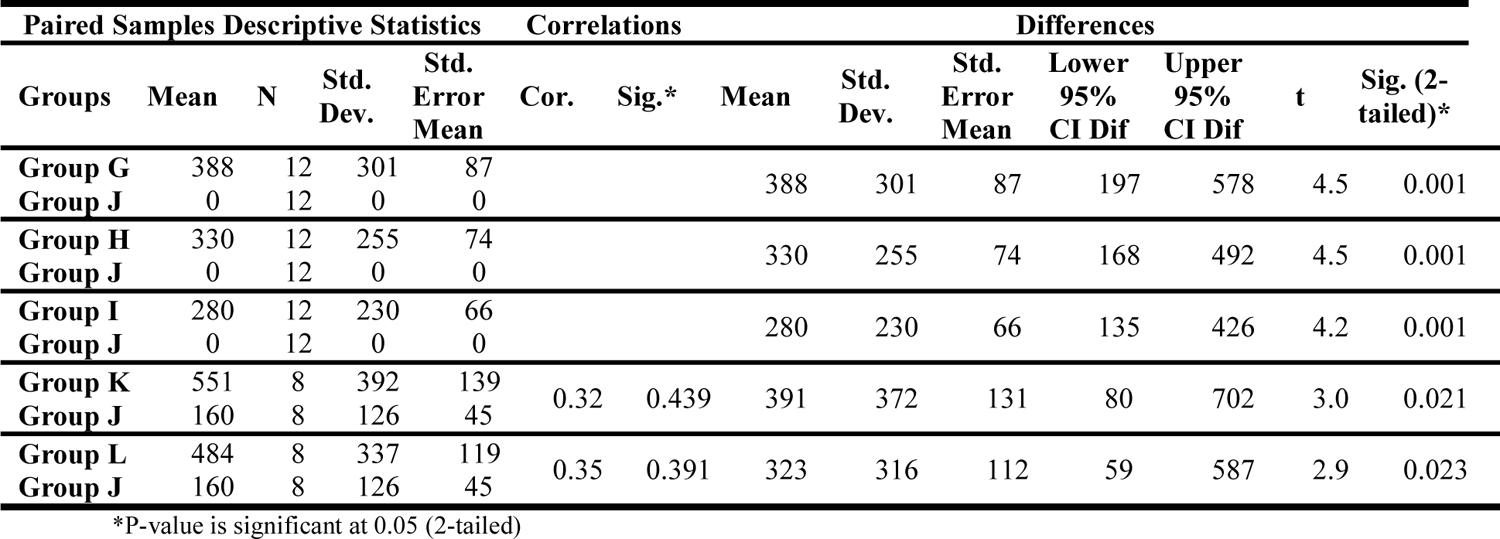
Paired t-test results of correlation and differences between vaccinated groups (G, H & I) and control group (J) as regards NAbs levels.

| Paired Samples Descriptive Statistics |  |  |  |  | Correlations |  | Differences |  |  |  |  |  |  |
| --- | --- | --- | --- | --- | --- | --- | --- | --- | --- | --- | --- | --- | --- |
| Groups | Mean | N | Std. Dev. | Std. Error Mean | Cor. | Sig.* | Mean | Std. Dev. | Std. Error Mean | Lower 95% CI Dif | Upper 95% CI Dif | t | Sig. (2-tailed)* |
| Group G | 388 | 12 | 301 | 87 |  |  | 388 | 301 | 87 | 197 | 578 | 4.5 | 0.001 |
| Group J | 0 | 12 | 0 | 0 |  |  |  |  |  |  |  |  |  |
| Group H | 330 | 12 | 255 | 74 |  |  | 330 | 255 | 74 | 168 | 492 | 4.5 | 0.001 |
| Group J | 0 | 12 | 0 | 0 |  |  |  |  |  |  |  |  |  |
| Group I | 280 | 12 | 230 | 66 |  |  | 280 | 230 | 66 | 135 | 426 | 4.2 | 0.001 |
| Group J | 0 | 12 | 0 | 0 |  |  |  |  |  |  |  |  |  |
| Group K | 551 | 8 | 392 | 139 | 0.32 | 0.439 | 391 | 372 | 131 | 80 | 702 | 3.0 | 0.021 |
| Group J | 160 | 8 | 126 | 45 |  |  |  |  |  |  |  |  |  |
| Group L | 484 | 8 | 337 | 119 | 0.35 | 0.391 | 323 | 316 | 112 | 59 | 587 | 2.9 | 0.023 |
| Group J | 160 | 8 | 126 | 45 |  |  |  |  |  |  |  |  |  |

\*P-value is significant at 0.05 (2-tailed)

To analyze the vaccine’s immunogenicity, the humoral and cellular immune responses were evaluated by different methods including qualitative IC-specific IgM and IgG-antibody titers (ELISA), virus-neutralizing assay in serum samples, biochemical and hematological analysis in peripheral blood samples, cytokines assay (western blotting), histopathology and immunohistochemistry in lung tissue. All inflammatory and immunological marker levels were lowered or became within normal ranges with advanced days post-immunization (Fig 3). In addition, the results of the Virus Neutralizing Test (VNT) in mice groups showed that the serum titer was 1/512. As regards inflammatory and immunological markers, statistical analysis by one-way ANOVA confirmed that there were highly significant differences between the CRCx3 groups (A, B & C) and the control group D but non-significant differences with the CRCx2 groups (E & F). The results of western blot showed that IL-6 decreased significantly in vaccinated animals compared to the control group (*P* < 0.05) (See Supplement; Table 2, Fig. 7).

Increasing serum anti-ICs antibody titers in vaccinated mice was monitored from the beginning and after days 14, 28, 42 & 56 (ELISA). The results showed that all of the vaccinated animals had significantly higher serum neutralizing antibody titers than the mock group (controls). Furthermore, the seroconversion rate in all vaccinated groups reached 100% by day 42 (Fig. 2). The CRCx3-vaccinated groups took a longer time to reach their maximum titers of NAbs than the CRCx2-vaccinated groups, which reached their maximum earlier. By day 42, CRCx3 groups had not reached their maximum and went on until after day 56 while CRCx2 groups reached their maximum by day 42 and then started declining to be less after day 56. This is probably because the first are nonspecific-ICs NAbs and the body immune system was taking its time to recognize the new antigenic combinations while the latter was faster to react with the body immune system being already of familiar structure. The highest produced NAb was Anti-B1 produced against the immune complex of the spike protein and its specific anti-spike antibody. CRCx3 high dose group A and CRCx2 high dose group E showed the highest mean titers with each vaccine, respectively. In addition, the mean NAbs titers for the double injection mode of CRCx2: [0 (B1)/7 (B2)] of HD (group E) and LD (group F), were higher than the triple injection mode of CRCx3: [0 (A1)/7 (A2)/14 (A3)] in HD (group A), MD (group B) and LD (group C); i.e. anti-specific ICs’ NAbs boosted more than anti-non-specific ICs’ NAbs in the sera of vaccinated mice (Fig. 1) (See Supplement: Table 3, Fig. 8). Statistical analysis by one-way ANOVA confirmed that on seroconversion by day 42, the 2-dose CRCx2 vaccines produced higher levels of NAbs Anti-B1 and Anti-B2 (Fig. 2) (See Supplement: Table 3, Fig. 8). A paired t-test between vaccinated groups and the control group in relation to NAbs showed highly significant differences between all vaccinated groups and the control group and showed highly significant correlations only in CRCx2 groups (E & F) with the control group (Table 1) (Fig. 1).

**Table 3:** Mean Rabbit body temperature (°C) after the pyrogenicity test.

| Rabbits Groups | Initial temperature | After 1 hour | After 2 hours | After 3 hours |
| --- | --- | --- | --- | --- |
| Group 1 | 38.3 | 38.5 | 38.6 | 38.6 |
| Group 2 | 38 | 38.2 | 38.4 | 38.4 |
| Group 3 | 38.5 | 38.6 | 38.7 | 38.7 |
| Group 4 | 38.4 | 38.6 | 38.6 | 38.5 |
| Group 5 | 38.1 | 38.2 | 38.4 | 38.6 |

To examine the disease severity and viral protein of SARS-CoV-2, the lungs in different groups were analyzed with H&E. Control group showed severe pathological changes denoting severe interstitial pneumonia including peribronchiolar and perivascular inflammatory infiltrations, alveolar wall thickening, and fibrin exudation. In contrast, the pathological changes were almost absent in the lungs of the CRCx2 high dose group and much milder in the CRCx2 low dose group showing normal lungs with focal mild histopathological changes in a few lobes with the CRCx3 groups. Generally, the average pathology score was lower in the vaccinated mice groups than in the placebo groups. All mice that received vaccination showed normal lung with focal mild histopathological changes in a few lobes. Collectively, the candidate vaccine was highly efficacious in mice at the two dosage levels with CRCx2 (Fig. 3).

### 2.2. Therapeutic Use in Mice

The mice in groups (G, H, I, J, K, and L) were challenged for 14 days by direct inoculation of 10^6^ TCID_50_ of SARS-CoV-2 intramuscularly. First, before vaccination with either CRCx, a few mice in every group died by day 12 dpi. All mice in groups showed marked elevation in temperature, lethargy, weight loss, and sneezing. They also developed ruffled fur, a hunched posture, and rapid breathing. Collectively, all mice showed signs of pneumonia with marked elevation in body temperature, nervous manifestations, and marked elevations in all biochemical and biological markers.

BALB/c mice were injected with triple combinations [15 (A1)/22 (A2)/29 (A3) days in groups G (High dose), H (Medium dose) & I (Low dose)] and double [15 (B1)/22 (B2) days in groups K (High dose) & L (Low dose)] immunization programs. In other words, the BALB/c mice in groups G & K received high (HD), H received medium (MD) and I & L received low (LD) doses of the candidate vaccines. Group J received normal saline as a placebo.

Throat and anal swabs that were obtained from mice on days 25 (first 4 groups) and 40 (last 2 groups) for viral load were measured by real-time PCR and confirmed higher viral load in the placebo group than in the challenge groups. On day 21 post-vaccination all vaccinated groups showed significant symptomatic improvement with a decrease in body temperature. By day 30, they showed a marked decrease in viral load to become normal as well as all the biochemical and immunological markers. For immunogenicity, the humoral and cellular immune responses were evaluated by different methods including qualitative IC-specific IgM and IgG-antibody titers (ELISA), virus-neutralizing assay in serum samples, biochemical, and hematological analysis in peripheral blood samples, cytokines assay (western blotting), histopathology and immunohistochemistry in lung tissue. Statistical analysis by one-way ANOVA revealed highly significant differences between all the vaccinated groups and the control group (J) with improvement in all indices for the vaccinated groups (See Supplement: Table 4, Fig. 9). In particular, IL-6 by western blot showed that IL-6 decreased significantly in vaccinated animals compared to the control group (*P* < 0.05).

Increasing serum anti-ICs antibody titers (ELISA) in mice was monitored after 14-day (dpi) on days 14, 28, 42 & 56 from the start of the experiment. Seroconversion rate in all vaccinated groups reached 80% by 28 days and 100% by day 42. On seroconversion on day 42, both anti-A1 and anti-B2 were produced more than other NAbs (Fig. 5). The results showed that all of the vaccinated animals had significantly higher serum neutralizing antibody titers than the mock (control) group. Moreover, the highest produced NAb was Anti-B1 produced against the immune complex of the spike protein and its specific anti-spike antibody. CRCx3 high dose group G and CRCx2 high dose group K showed the highest mean titers with each vaccine, respectively. Even though both candidate vaccines generally induced high levels of NAbs, the mean NAbs titers for the double injection mode of CRCx2: [15 (B1)/22 (B2)] of HD (group K) and LD (group L), were higher than the triple injection mode of CRCx3: [15 (A1)/22 (A2)/29 (A3)] in HD (group G), MD (group H) and LD (group I); i.e. anti-specific ICs’ NAbs boosted more than anti-non-specific ICs’ NAbs in the sera of vaccinated mice (Fig. 4) (See Supplement: Table 5, Fig. 10). Statistical analysis by a paired t-test between groups and NAbs showed significant differences between all vaccinated groups and the control group (J). A paired t-test between the vaccinated groups and the control group with NAbs showed highly significant differences between all vaccinated groups and the control group (Table 2).

### 2.3. Evaluation of (CRCx2) and (CRCx3) Safety

In this study, 24 mixed-gender rats were divided into four groups, one control, and three treatment groups. The rats in the three treatment groups were intramuscularly injected with two doses (Triple High=1.2 ml, Triple Medium=0.8 ml, or Triple Low=0.6 ml) of candidate vaccines and physiological saline as the control on days 0 and 7. All rats were monitored for 14 days to assure that there were no cases of death, any significant difference in weight or feeding state, or any histopathologic changes after euthanasia (15 days after vaccination). Results showed no cases of death, impending death, or obvious clinical signs including local injection irritation in any of the treatment groups over 14 consecutive days. Moreover, there was no significant difference in weight or feeding state between the experimental groups and the control group. In addition, no significant histopathologic changes were observed after euthanasia (Fig. 6). It should be noted that the maximum tolerated dose (MTD) used for a “triple high” shot in rats (1.2 ml/rat) is six times the dose in humans, which makes it a reliable candidate vaccine as a matter of safety, indicating the potential good safety of CRCx in humans. Monitoring lymphocyte subgroup distribution (CD4+, CD8+ and the CD4+/CD8+ ratio) in all tested animals showed that neither CD4+, the CD8+ nor the CD4+/CD8+ ratio, differed after vaccination. This finding was irrespective of either immunization program or administrated dosage (Fig. 7).

The results of the pyrogenicity assay in rabbits (Table 3) showed that the total increase in body temperature of the five rabbits groups at 3 different intervals was below the threshold (>1.2°C). Also, the difference between the body temperature of each animal at zero hours, and the following hours was up to 0.4°C. Therefore, it can be concluded that the candidate vaccine are not pyrogenic. Furthermore, systemic anaphylaxis due to CRCx was evaluated by intramuscular and intravenous injections in twenty-four guinea pigs showing high tolerance to CRCx dosage without acute single dose, multiple doses, or local injection irritation. In addition, the experiment revealed that the inactivation steps were independently capable of inactivating vaccine titers with a large margin of safety.

### 1.3. Discussion

The novel COVID-19 has demonstrated a rapid spread since December 2019 causing a huge outbreak in China [50]. The development of preventive and therapeutic vaccines with high immunogenicity and safety is crucial for the control of the global COVID-19 pandemic and the prevention of further morbidities and fatalities [51]. Accordingly, scientists have focused their efforts on the development of vaccines to stop the COVID-19 pandemic and its quick spread [52]. The first genome sequence of SARS-CoV-2 was reported in China on 11 January 2020, encouraging exceptional efforts to develop various vaccine candidates [53]. Until a few months ago, there were 123 candidate vaccines in human clinical trials and 194 candidates in preclinical development worldwide [11].

One of the main concerns in the development of vaccines is the antibody-dependent enhancement of coronavirus disease phenomenon, better known as ADE [54], which whereby the vaccine could make the subsequent SARS-CoV-2 infection more severe [55] & [56]. ADE mechanisms involve immune complexes formed between the virus and non-neutralizing antibodies and poorly neutralizing antibodies that bind to Fcy receptors (FcyRs), which are expressed broadly on monocytes and macrophages leading to the internalization of the virus particle to enter the cell [57]. The ADE phenomenon may happen due to an imbalance between the T-helper 1 and T-helper 2 responses. Granzyme B, a neutral serine protease, is expressed exclusively in the granules of activated cytotoxic T-lymphocyte, natural killer cells, and lymphokine-activated killer cells. Granzyme B induces T-lymphocyte-mediated DNA fragmentation and apoptosis in allogeneic target cells [58]. Previous studies noted the importance of the immune complexes as inflammatory mediator stimuli. An immune complex is formed from a neutralizing Ag/Abs complex [59], [60]. Normally, the insoluble immune complexes that are formed are cleared by the phagocytic cells of the immune system, but when an excess of antigens/antibodies is present, the immune complexes are often deposited in tissues, where they can elicit complement activation and localized inflammation, resulting in the generation of tissue lesions in a variety of autoimmune diseases [61], [62], [63], which exacerbates the disease pathology.

In a previous research we had conducted on AIDS, we concluded that there is an alteration in the physiological behavior of CD4+ T-cells causing it to mutate to CD8+ T-cells and that CD4+ T-cells are neither destroyed nor lost during the infection [64]. Similarly, we think that instead of helping to eliminate circulating SARS-CoV-2, circulating non-neutralizing antibodies can bind to viral particles and thus, contribute to the worsening of COVID-19 by promoting its Fc-mediated internalization by pulmonary epithelial cells and infiltrating monocytes as was previously observed in SARS-CoV-1 [54] and SARS-CoV vaccines in animal challenge models [64]. Covid-19 virus, despite its external spikes that enable it to adhere to the cell surface, cannot determine the suitable adherence point and needs a familiar immune cell (CD4+ T-helper cell) to direct it to the appropriate tropism and link it to this receiving cell and, thus, the virus can live masquerading within our immune defenses as a biological part similar to our cells, giving it the advantage over our immune system. Once infected, the CD4+ T-helper cells will be controlled by the virus accepting the information embedded within the virus code leading to many immune and cellular changes. To start with, the infected cell will exchange with the virus the information to use to attack the respiratory cells as the second target cell and replicate. Then, the infected T-helper cell will send signals to excite the B-cells to produce a number of immune antibodies to attach to the viral antigens that are configured by the respiratory cells, thus forming a circulating immune complex that comprises coronavirus antigen (M, N, and S and its specific neutralizing antibody as IC_1_, IC_2_, and IC_3_). These formed circulating immune complexes will protect the virus from the attack of CD8+ killer cells. Afterward, the infected cell will infect other CD4+ T-cells granting the virus a continuation of the infection cycle. The first infected helper cells will become a modulated version of CD8+ T-cells but with different physiological behavior, functions, and divergence from the original CD4+ T-cells. The loss of CD4+ T-cells causes complete cellular chaos and discrepancies in the cell signals, e.g. the newly formed CD8+ T-cells may permit an activator signal while the original CD8+ T-cells inhibit it. This imbalance leaves the immune system in a state of confusion with two contradictory responses that directly cause dysregulation and ultimate failure of the host cellular immune networks (See Supplement: Fig. 1).

Neutralizing antibodies play a critical role in protection against SARS-CoV-2 infection and have been used as an immune correlate of protection in assessing vaccine efficacy. According to our postulation, the broadly neutralizing antibodies that are generated during vaccination exist in two forms. The first is a positive form that is bound to and effectively masks a proportion of the coronavirus antigen’s structural and nonstructural protein in a complex form that prevents its elimination. The second is a free negative form, which is nonfunctional and non-neutralizing. This immune complex formation can explain the persistence of the viral infection. Even with all approved vaccines worldwide, we cannot rule out the possibility that the evolution of the virus can still directly affect its targets, and therefore, the newly mutated virus can escape antibody-mediated protection induced by previous infection or vaccination. If the vaccines cannot generate enough neutralizing antibodies against the possible mutagenic variants to mount a response, the result might be the production of sub-neutralizing antibodies that will even be capable of facilitating uptake by macrophages expressing antibody Fc region reaction with cellular Fc receptors (FcRs), with the subsequent stimulation of macrophages and production of pro-inflammatory cytokines [54].

In our present study, we report the creation of a novel method for the development of a SARS-CoV-2 vaccine. CRCx was formulated with twenty-five micrograms (25 μg) of different antigens including spike protein (S1 subunit), nucleocapsid (N), or membrane antigen (M) as well as forty micrograms (40 μg) of different antibodies including anti-nucleocapsid, anti-membrane or anti-spike (S1 subunit) *antibodies.* Doses were mixed with Fc of IgG mouse anti-Human IgM added as an adjuvant immunogen dissolved in human albumin, phosphate buffer, and sodium chloride without any stabilizers or preservatives. The main aim of this novel immune peptide CRCx is to stimulate the CD8+ T-cells against the foreign antigenic non-complexes as well as any other hidden circulating immune complex form with any antigenic similarity (antigen/specific-antibodies) and to put the whole immunity system on alert and restoring its normal functions. The CD4+ T-cells will start sending a new signal that will interfere with the viral signals, stopping its antigens production in the respiratory cells as well as stopping the production of antibodies from B-cells and cessation of their infection of other CD4+ T-cells or mutation to physiologically unfamiliar CD8+ T-cell. Accordingly, the virus cycle and its operations are disrupted. In this way, the vaccine does not only boost immunity but also trains it to remember the virus if any modifications in its structure occur by frequent mutations.

By incorporating the different combinations of antigen/non-specific-antibody as well as antigen/specific-antibody formulas with an immunogenic adjutant (to facilitate response and recognition), the immunity system is rendered capable of recognizing and committing to memory all kinds of different antigenic combinations for future recognition. The initial experiments were designed to find out which is the best combination or formula that will produce the highest immunogenicity and highest neutralizing antibody titers in response to SARS-Cov-2 infection. Animal models that could adequately simulate the viral infection and its development similar to that in humans are regarded as a perfect choice to investigate the performance of vaccines. Since mice are a model of SARS-CoV infection but not disease so, Balb/C mice were used in the evaluation of developed SARS-COV2 vaccines [65]. Young BALB/c mice have higher antigen-binding and neutralizing antibody levels than those of old BALB/c mice.

In this study, we first analyzed the immunogenicity of the CRCx in young BALB/c mice. The CRCx formulations were found to develop high titers of neutralizing antibodies by mechanisms that are obscured. They may benefit the antibody reaction by favoring the activation and trafficking of antigen-presenting cells to lymphoid tissues in addition to triggering the inflammasome and complementing activation. Also, by provoking the CD8+ killer cells to recognize these circulating immune complexes created by the influence of the virus on CD4+ cells, CD8+ cells will treat them as foreign agents by shifting the immune system to an alert state, the immune cells’ normal physiological behavior can be restored.

As protective or prophylactic vaccines, our results show that the candidate vaccine formulations induced significantly elevated neutralizing antibody responses in the vaccinated mice with a high dose of CRCx2 and CRCx3 vaccines. However, the NAbs means were highest with the CRCx2 that were formulated of spike and nucleocapsid antigens and their specific antibodies, protecting mice against SARS-CoV-2 infection. The highest level of NAb produced in all groups was the Anti-B1, the one against spike/anti-spike complex. Similarly, as therapeutic vaccines, our results show that the candidate vaccine formulations induced significantly elevated antigen neutralizing antibody responses in the 14-28 days post-challenged vaccinated mice with similar results to the prophylactic approach but with higher titers of NAbs produced against all vaccine combinations. Fc of IgG mouse anti-Human IgM, which is used here for the first time as a vaccine adjuvant, proved an unparalleled safety record desired to obtain a COVID-19 vaccine that can generate both humoral and cell-mediated immune responses. In addition, strong neutralizing antibody production by different vaccine combinations was shown to be time-dependent and dose-dependent.

In addition, toxicity and safety evaluation showed no adverse or clinical signs in vaccinated mice. The immune-complex vaccine would be most efficient in inducing neutralizing antibodies, which are possibly critical in preventing SARS-CoV infection. The safety evaluation of CRCx showed no local or systemic toxic manifestations in mice, as demonstrated by the repeated dose toxicity with no changes in body weight, body temperature, or any other obvious clinical signs. Furthermore, there were no cases of death, impending death, or *post-mortem* histopathological changes. The maximum tolerated dose (MTD) used for a single intramuscular injection in rats was 1.2 ml/rat (equivalent to 6 times the dose in humans) indicating potentially good safety margins. The results of the pyrogenicity assay in rabbits concluded that the candidate vaccines are not pyrogenic. Systemic anaphylaxis due to CRCx was evaluated by intramuscular and intravenous injections in guinea pigs. The results showed no abnormal reactions during the sensitization period nor any allergic reaction symptoms in either experimental or control groups.

Finally, our data demonstrate complete protection both as a preventive vaccine and as an antiviral therapy against SARS-CoV-2 by inhibiting virus replication in lung tissue after a challenge test in the lung tissues of mice that received the vaccine. These data were confirmed by the absence of histopathological findings in the lungs of vaccinated mice groups. The results also did not find the vaccine to cause antibody-dependent enhancement (ADE) [66] as all the data obtained in this study support the safety and immunogenicity of this candidate vaccine series. The antigen/nonspecific-antibody-IC vaccine series needs more experiments to validate our primary hypothesis that they may prompt more potent and more long-lasting protection against mutating versions of SARS-Cov-2 as results showed that within the time limit of this trial, the antigen/specific-antibody-IC vaccine series produced higher NAbs against the vaccine ICs.

### 1.4. Conclusion

The new immune complex (IC) anti-SARS-Cov-2 candidate vaccine CRCx is composed of 5 different combinations: a series of three antigen/nonspecific-antibody termed (CRCx3) and two antigen/specific-antibody termed (CRCx2). Our animal model trial proved that both CRCx2 and CRCx3 vaccine candidates can induce highly efficient protection and therapeutic efficacy against SARS-CoV-2 without observable antibody-dependent enhancement (ADE) or immune-pathological exacerbations. Generally, in both prophylactic and therapeutic approaches trials, the highest neutralizing antibody (NAb) produced in all groups was the anti-B1, which is the NAb against the vaccine composed of spike antigen and its specific antibody of anti-spike. Specifically, the highest titer of this NAb was seen in the groups that were immunized with high doses of this vaccine combination which is a CRCx2 sub-type. In addition, due to the trial time limit of 56 days, the groups immunized with CRCx3 subtype combinations did not have enough time to reach their maximum production of corresponding NAbs. However, again, the highest titer in these groups was seen with the high dose CRCx3 subtype composed of spike and its nonspecific antibody of anti-nucleocapsid.

Conclusively, according to the results, the spike antigen/anti-spike specific-antibody combination of CRCx2 gives the highest immunogenicity against Covid-19 virus infection both as a prophylaxis and as a therapy. In addition, even though both vaccines were found to significantly reduce or abolish viral load and broncho-alveolar effects in animal models challenged with SARS-CoV-2 within 14 days after receiving the booster dose of the vaccines with no signs of pneumonia in histopathological sections of the virus-challenged animals after vaccination, a higher preference was found to the double-dose antigen-specific-antibody (CRCx2) series and/or those from both CRCx series with the spike protein antigen. As for the toxicity of the vaccine series, single-dose and repeated-dose toxicity showed no abnormal changes related to vaccination in animals when given a triple high dosage of vaccine alone to otherwise healthy control animals. As for active systemic anaphylaxis, no allergic reaction was observed after the guinea pigs were injected with a high dosage vaccine intended for clinical use. Furthermore, candidate vaccines are not pyrogenic. Finally, in the absence of an effective antiviral drug against SARS-CoV-2, vaccines with appreciable potency and safety are needed to effectively establish immunity in the population. Based on the results presented in this pretrial, a phase I clinical trial of CRCx is recommended to be undertaken.

## Author Contributions

Authors’ Contributions are not equal. S. Salah conceived the study; S. Sherif, M. Abdula & Z Khalid designed the experiments. M Abdula performed the experiments. Z. Khalid analyzed the data and wrote the manuscript. O. Khaled reviewed and edited the manuscript.

## Institutional Review Board Statement

The study was conducted according to the guidelines of the Declaration of Helsinki and approved by the Committee for Ethics on Animal Experiments and the Committee for Animal Biosafety Level 3 Research of the Egyptian Military Scientific Commission.

## Informed Consent Statement

All authors have read and agreed to the published version of the manuscript.

## Data Availability Statement

The data that support the findings of this study are available on request from the corresponding author.

## SUPPLEMENT

**Figure 8:**
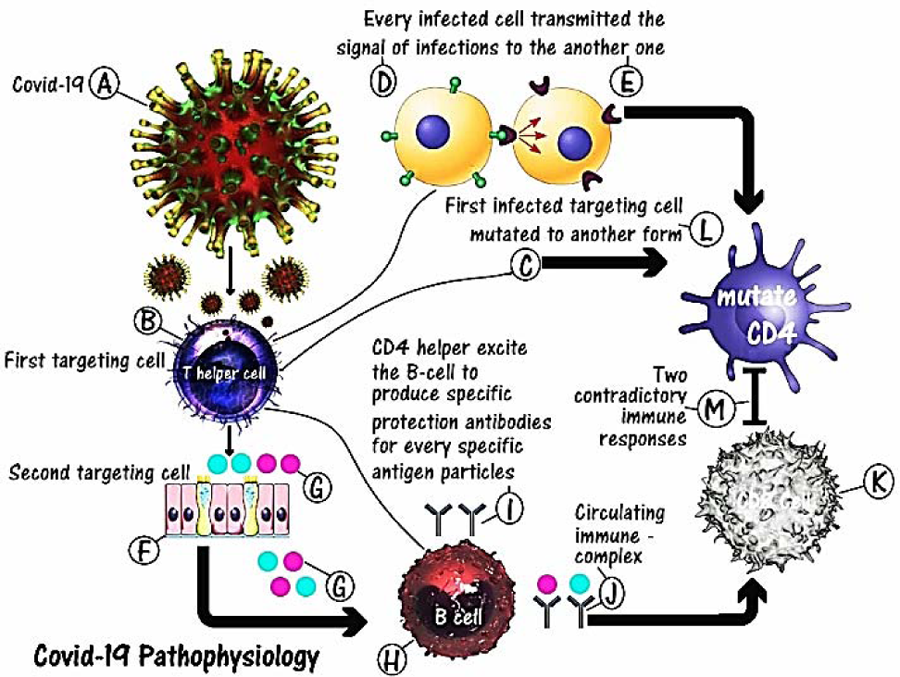
Hypothetical model of action for Coronavirus. (A) Virus particles. (B) CD4+ T-cell as the first susceptible cell. (C) Evolution of the CD4+ T-cell mutating to be an unaccountable CD8+ T-cell. (D) Tendencies of the first susceptible infected cell to infect other healthy ones and to be a new form of CD8+ T-cells. (E) Newly formed infected first susceptible cell. (F) Cells of the upper and lower respiratory tract as second susceptible cells. (G) Productions of viral antigen particles from the second susceptible cells under the induction of the first susceptible cells’ stimulus of transmitted ions signals. (H) CD4+ T-cell stimulating B-cells to produce negative neutralized coating antibodies to form a complex with antigen particles. (I) B-cells produce negative neutralizing antibodies. (J) Formations of circulating immune complex comprising coronavirus antigens and its specific negative neutralizing antibodies. (K) CD8+ T-cells. (L) Inhibitory effect of these complexes in preventing the cytotoxic CD8+ T-cells from attacking these viral particles. (M) The interaction mechanisms that originate as a result of antagonism between the newly formed mutated CD4+ T-cells and the normal CD8+ T-cells.

**Figure 9:**
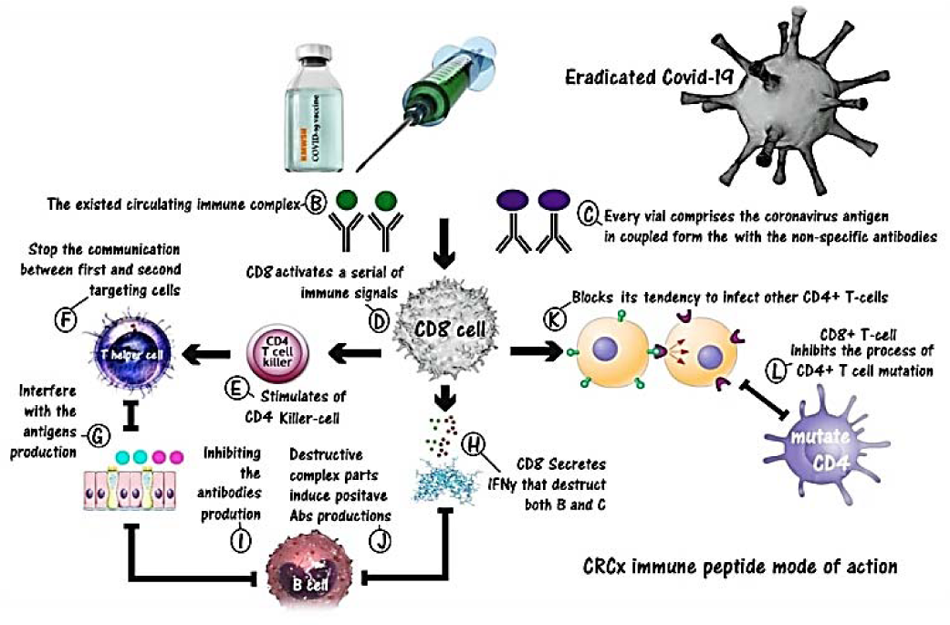
Immunity stimulating action of the CRCx3 vaccine. (A) CRCx immune peptide. (B) Existing circulating immune complex. (C) Every vial comprises the coronavirus antigen in coupled form with non-specific antibodies. (D) CD8+ T-cell stimulated to induce a process of scanning and comparison between these non-complex combinations and those already existing during the vaccine injection. Also. CD8 stimulates a series of immune signals. (E) Stimulation of CD4+ killer cells. (F) Stimulation of CD4+ Helper cell. (G) CD4+ T-cells send a signal that inhibits the formations of the coronavirus antigens. (H) Cytotoxic T-cells secreting IFNy that destroys the circulating immune complex and the intruder non-complexes. (J) Destructive complex particles induce positive antibody productions from B-cells. (K) Blocking of the tendency of CD4+ to infect other cells. (L) CD8+ T-cells inhibit the process of CD4+ T-cell mutation.

**Figure 10:**
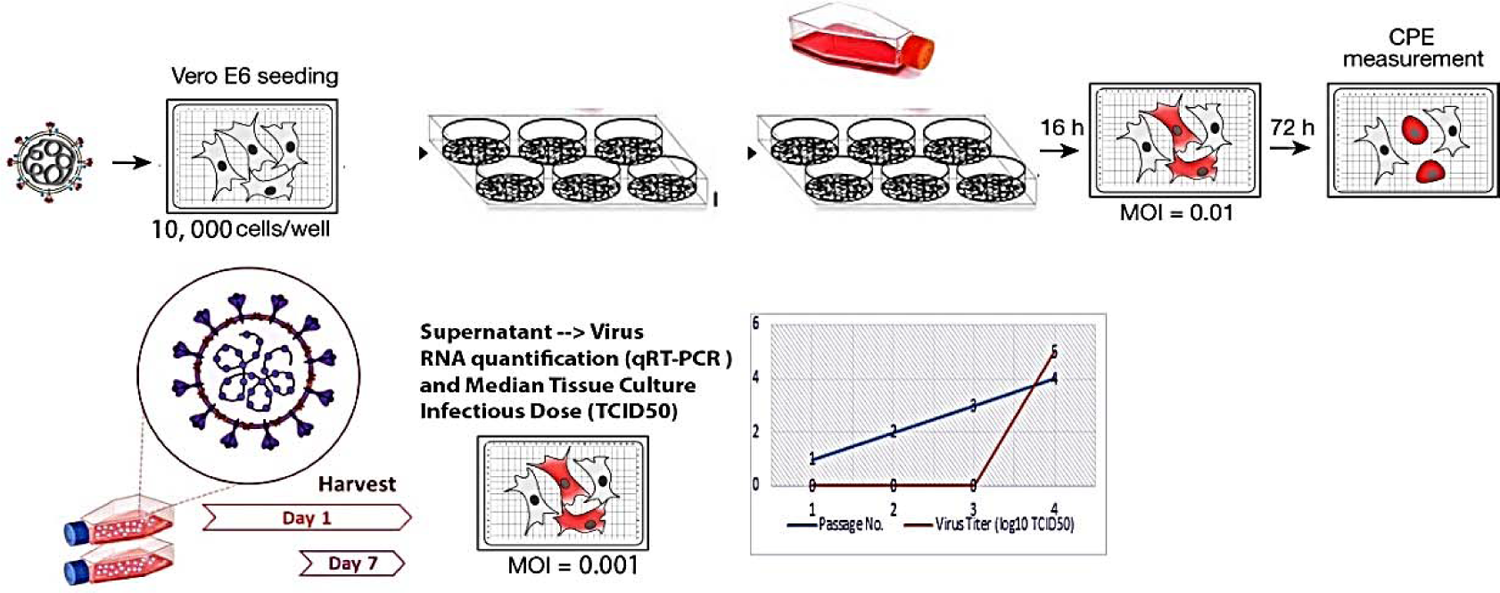
Preparation and titration of SARS-CoV-2 Stock Solution

**Figure 11:**
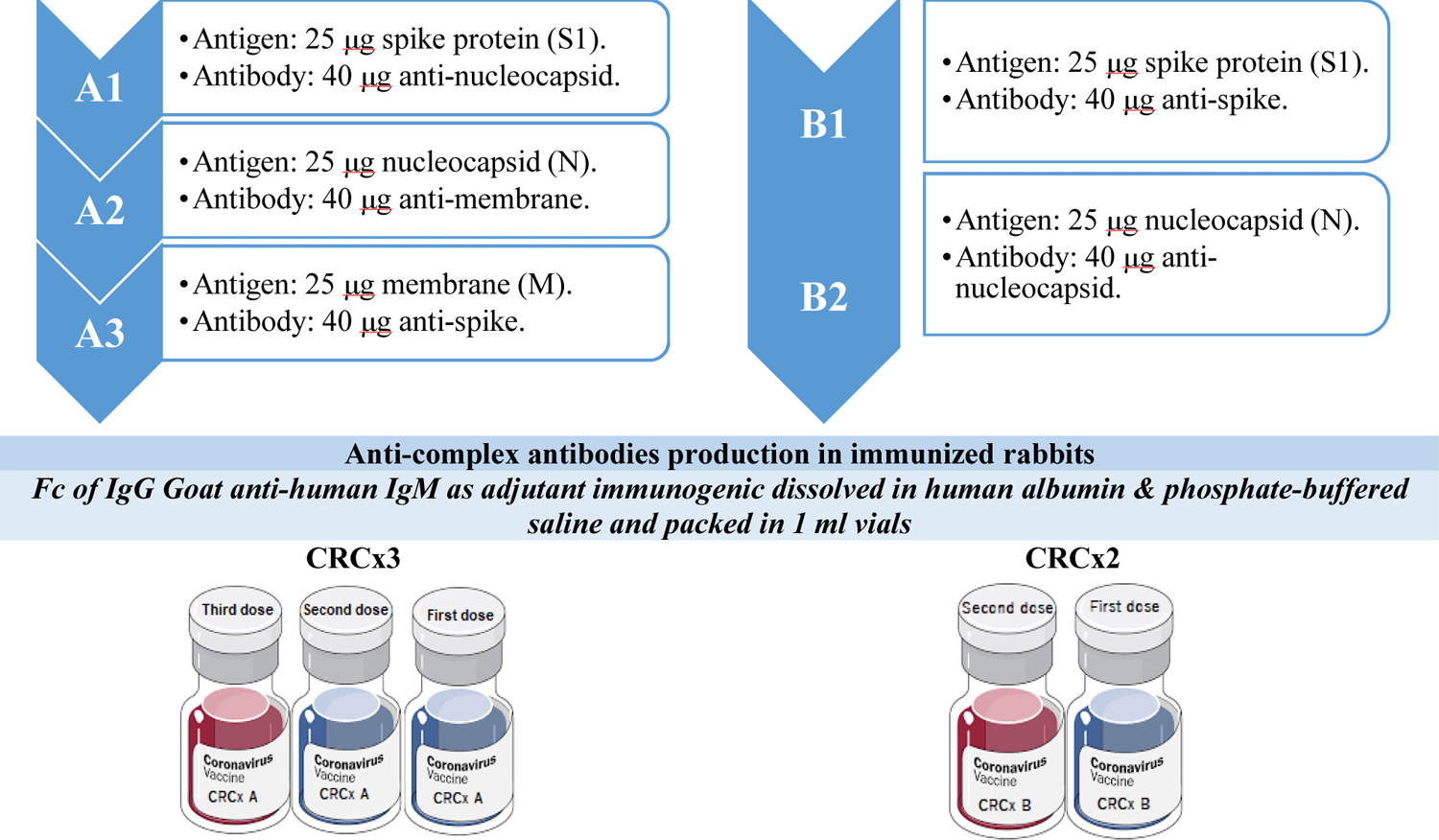
Formulation of CRCx2 & CRCx3

**Figure 12:**
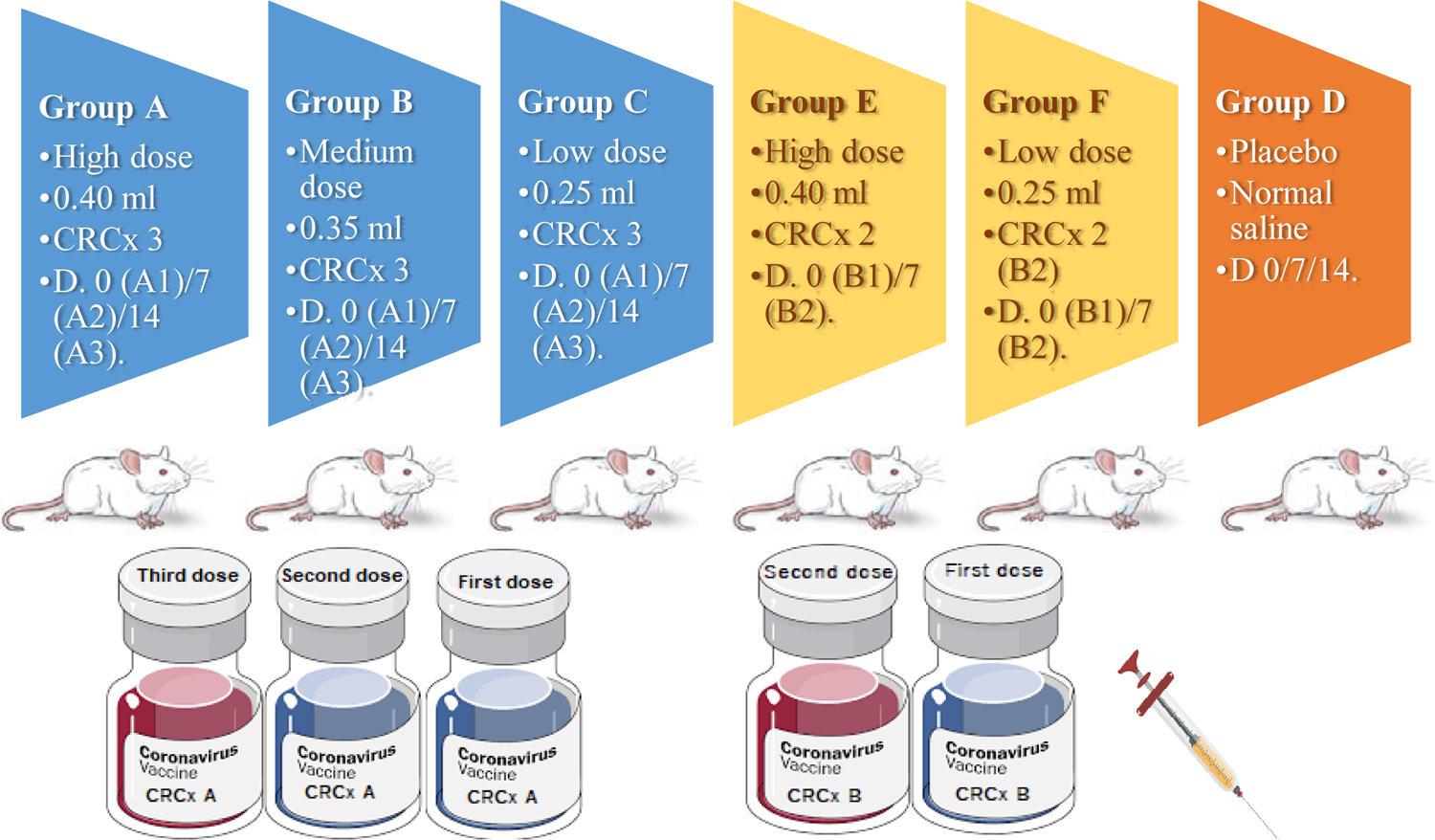
CRCx vaccines experiment as prophylactic candidates in Balb/c mice.

**Figure 13:**
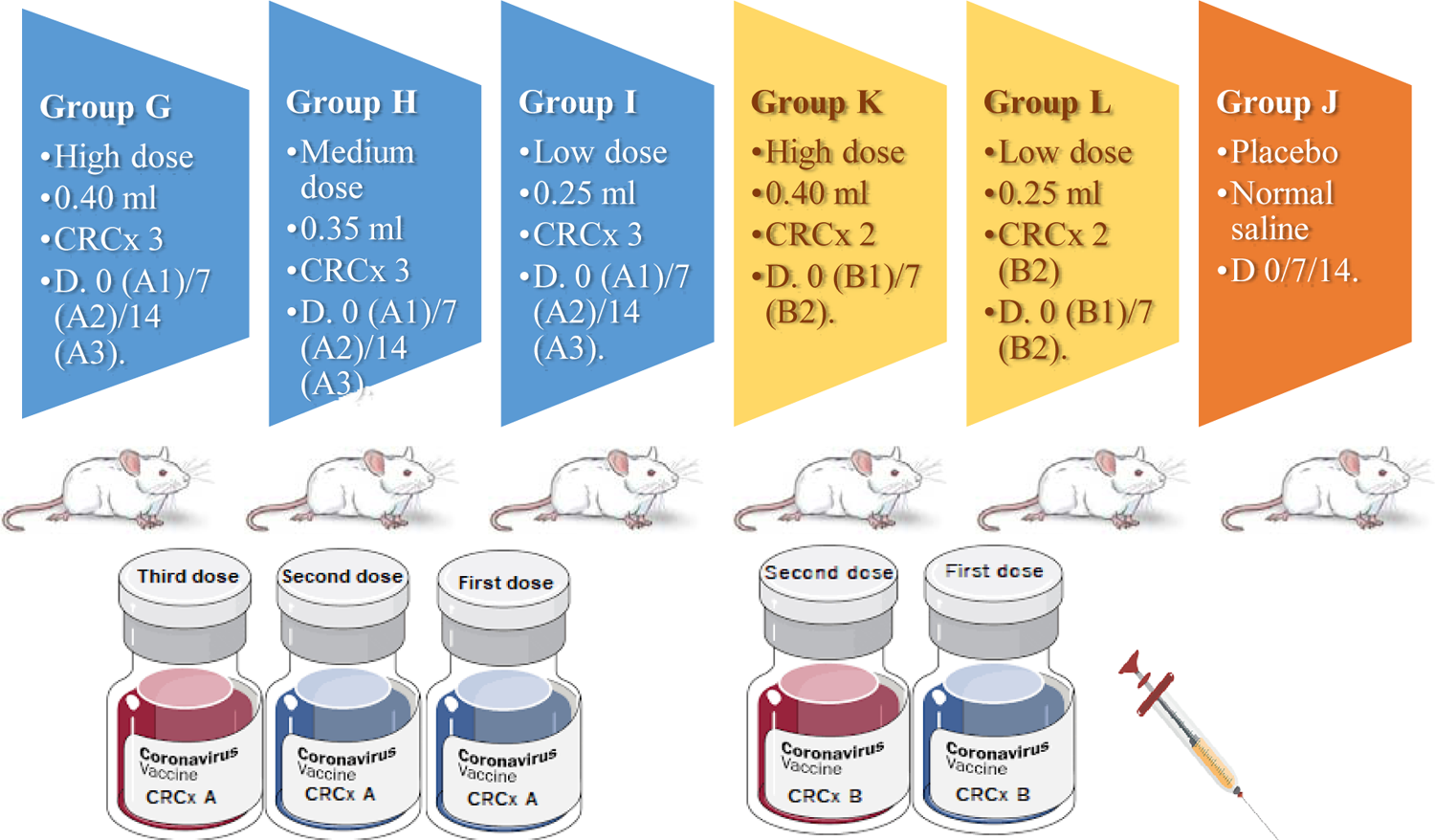
CRCx vaccines experiment as therapeutic candidates in Balb/c mice.

**Table 4:** Animal studies for evaluating the safety and efficacy of the CRCx series vaccine.

| Study | Topic | Animal model |
| --- | --- | --- |
| **Safety Assay** | Abnormal toxicity test / Single dose toxicity/ Acute toxicity | Rats |
|  | Repeated dose toxicity | Rats, Guinea pigs |
|  | Pyrogenicity assay | Rabbits |
|  | Active systemic anaphylaxis | Guinea pigs |
|  | Local injection irritation | Mice, Rats, Guinea pigs, Rabbits |
| **Efficacy: Vaccine immunogenicity analysis (Prophylactic/Therapeutic)** | Evaluation of the humoral immune response (IgG antibody titer 'ELISA'), Blood biochemical and hematological analysis, Virus Neutralizing Test (VNT), Cytokines assay | Mice, Rabbits |
|  | Evaluation of the cellular immune response ( SARS-CoV-2 challenge): Virus quantification in tissues, Cytokines assay, inflammatory markers, Histopathology and immune-histochemistry | Mice, Rats |

**Table 5:** Immuno-chemistry studies at 14, 28, 42 & 56 days after immunization with CRCx candidate vaccines in prophylactic trails mice groups.

| Animal Groups | Days | Group A | Group B | Group C | Group D | Group E | Group F |
| --- | --- | --- | --- | --- | --- | --- | --- |
| CD4+<br>(Cells/μl) | 14 Days | 285 | 305 | 321 | 1550 | 550 | 420 |
|  | 28 Days | 220 | 180 | 140 | 430 | 330 | 310 |
|  | 42 Days | 550 | 510 | 470 | 280 | 820 | 780 |
|  | 56 Days | 790 | 670 | 600 | 120 | 1300 | 1230 |
|  | Normal Range | 359-1954 |  |  |  |  |  |
| CD8+<br>(Cells/μl) | 14 Days | 720 | 600 | 540 | 900 | 830 | 770 |
|  | 28 Days | 930 | 820 | 760 | 850 | 650 | 550 |
|  | 42 Days | 520 | 600 | 500 | 560 | 440 | 340 |
|  | 56 Days | 390 | 430 | 400 | 400 | 400 | 320 |
|  | Normal Range | 220-1100 |  |  |  |  |  |
| CRP<br>(ng/ml) | 14 Days | 6 | 5.3 | 3.8 | 0.8 | 8 | 11 |
|  | 28 Days | 18 | 24 | 30 | 120 | 10 | 13 |
|  | 42 Days | 42 | 49 | 55 | 45 | 60 | 55 |
|  | 56 Days | 20 | 22 | 27 | 40 | 14 | 22 |
|  | Normal Range | 0.78-25 |  |  |  |  |  |
| D-dimer<br>(ng/ml) | 14 Days | 58 | 55 | 50 | 0.7 | 22 | 15 |
|  | 28 Days | 58 | 61 | 66 | 65 | 54 | 58 |
|  | 42 Days | 34 | 37 | 42 | 44 | 19 | 22 |
|  | 56 Days | 17 | 22 | 28 | 42 | 15 | 14 |
|  | Normal Range | 0.6-50 |  |  |  |  |  |
| IL-6<br>(ng/ml) | 14 Days | 0.8 | 0.5 | 0.4 | 1.4 | 0.4 | 0.2 |
|  | 28 Days | 1.6 | 1.9 | 2.4 | 2.2 | 1.2 | 1.4 |
|  | 42 Days | 0.8 | 1.2 | 1.3 | 3 | 0.16 | 0.13 |
|  | 56 Days | 0.15 | 0.33 | 0.4 | 14 | 0.03 | 0.06 |
|  | Normal Range | 0.01-0.2 |  |  |  |  |  |
| Qualitative<br>IgM | 14 Days | Negative | Negative | Negative | Negative | Negative | Negative |
|  | 28 Days | Positive | Positive | Positive | Positive | Positive | Positive |
|  | 42 Days | Positive | Positive | Positive | Positive | Positive | Positive |
|  | 56 Days | Positive | Positive | Positive | Positive | Positive | Positive |
| Qualitative<br>IgG | 14 Days | Negative | Negative | Negative | Negative | Negative | Negative |
|  | 28 Days | Negative | Negative | Negative | Negative | Negative | Positive |
|  | 42 Days | Positive | Positive | Positive | Negative | Positive | Positive |
|  | 56 Days | Positive | Positive | Negative | Positive | Positive | Positive |
| Qualitative<br>RNA | 14 Days | Negative | Negative | Negative | Negative | Negative | Negative |
|  | 28 Days | Positive | Positive | Positive | Positive | Positive | Positive |
|  | 42 Days | Positive | Positive | Negative | Positive | Positive | Positive |
|  | 56 Days | Negative | Negative | Negative | Positive | Negative | Negative |

**Figure 14:**
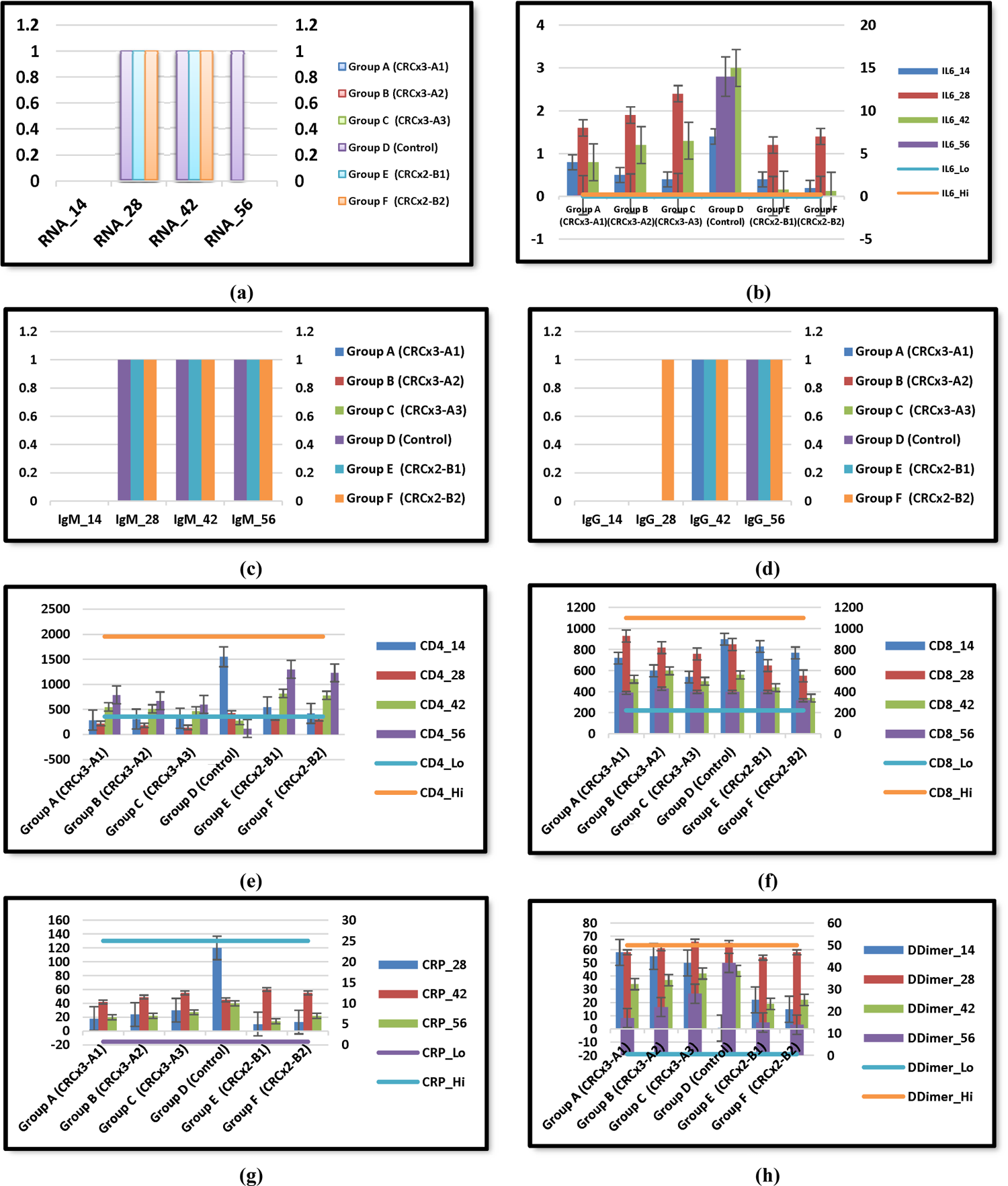
Quantitative and Qualitative Immunological and Biochemical Assays for Prophylactic Approach Groups on Days 14, 28, 42 & 56 with CRCx3 (Groups A, B & C), CRCx2 (Groups (E & F) and Control (D): (a) Qualitative SARS-Cov-2 RNA. (b) Quantitative IL 6. (c) Qualitative SARS-Cov-2 IgM. (d) Qualitative SARS-Cov-2 IgG. (e) Quantitative CD4+. (f) Quantitative CD8+. (g) Quantitative C-Reactive Proteins (CRP). (h) Quantitative D-dimer. Note: In quantitative assays, the horizontal lines represent the low and high normal values. In qualitative assays, 1=Positive; 0=Negative.

**Table 6:** Neutralizing antibodies (NAbs) titers on days 14, 28, 42 and 56 in prophylactic regimens.

| Animal Groups | Days | Group A | Group B | Group C | Group D | Group E | Group F |
| --- | --- | --- | --- | --- | --- | --- | --- |
|  | Regimen | CRCx3<br>(0.4x3) +<br>Cov-2 | CRCx3<br>(0.35x3) +<br>Cov-2 | CRCx3<br>(0.25x3) +<br>Cov-2 | NaCl + Cov | CRCx2<br>(0.4x2) +<br>Cov-2 | CRCx2<br>(0.25x2) +<br>Cov-2 |
| Anti-A1<br>(ng/ml) | 14 Days | 200 | 110 | 67 | 0 |  |  |
|  | 28 Days | 550 | 310 | 200 | 0 |  |  |
|  | 42 Days | 650 | 520 | 320 | 0 |  |  |
|  | 56 Days | 340 | 300 | 180 | 0 |  |  |
|  | Normal Range |  |  | 0.2-1 |  |  |  |
| Anti-A2<br>(ng/ml) | 14 Days | 85 | 72 | 57 | 0 |  |  |
|  | 28 Days | 260 | 210 | 170 | 0 |  |  |
|  | 42 Days | 430 | 300 | 240 | 0 |  |  |
|  | 56 Days | 150 | 120 | 90 | 0 |  |  |
|  | Normal Range |  |  | 0.5-1 |  |  |  |
| Anti-A3<br>(ng/ml) | 14 Days | 10 |  | 0 | 0 |  |  |
|  | 28 Days | 250 | 200 | 150 | 0 |  |  |
|  | 42 Days | 600 | 480 | 350 | 0 |  |  |
|  | 56 Days | 410 | 340 | 300 | 0 |  |  |
|  | Normal Range |  |  | 0.5-5 |  |  |  |
| Anti-B1<br>(ng/ml) | 14 Days |  |  |  | 0 | 90 | 75 |
|  | 28 Days |  |  |  | 5 | 450 | 410 |
|  | 42 Days |  |  |  | 180 | 800 | 780 |
|  | 56 Days |  |  |  | 240 | 600 | 560 |
|  | Normal Range |  |  |  | 0.5-2 |  |  |
| Anti-B2<br>(ng/ml) | 14 Days |  |  |  | 0 | 10 | 6 |
|  | 28 Days |  |  |  | 2 | 230 | 180 |
|  | 42 Days |  |  |  | 80 | 630 | 500 |
|  | 56 Days |  |  |  | 95 | 450 | 400 |
|  | Normal Range |  |  |  | 0.2-2 |  |  |

**Figure 15:**
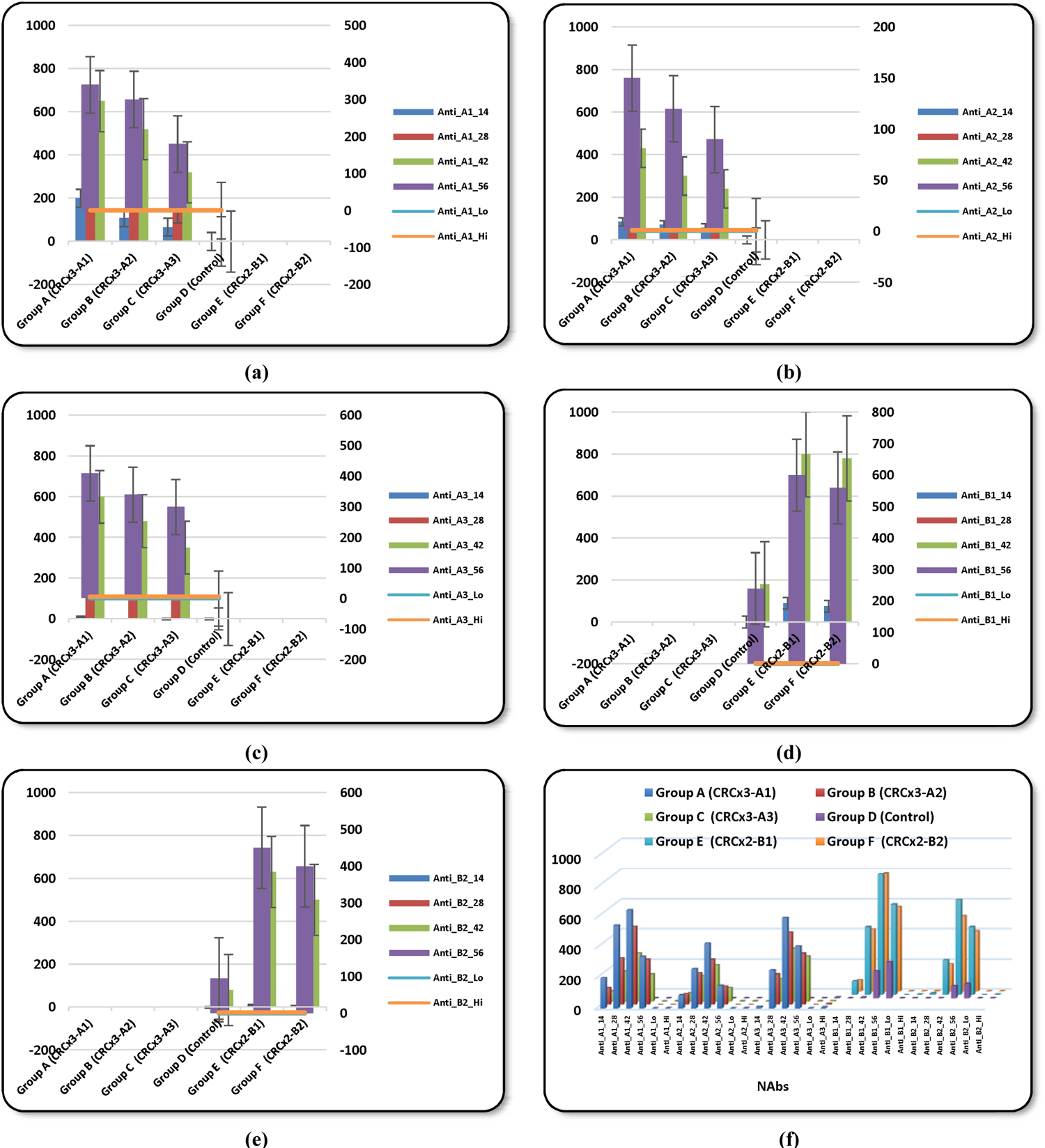
Different neutralizing antibodies (NAbs) in Prophylactic Approach Groups on Days 14, 28, 42 & 56 with CRCx3 (Group A, B & C), CRCx2 (Groups (E & F), and Control (D): (a) Anti-A1. (b) Anti-A2. (c) Anti-A3. (d) Anti-B1. (e) Anti-B2. (f) All NAbs. Note: The horizontal lines represent the low and high normal values.

**Table 7:** Immuno-chemistry studies at 42 & 56 days after immunization with CRCx candidate vaccines in therapeutic trails mice groups.

| Animal Groups | Days | Group G | Group H | Group I | Group J | Group K | Group L |
| --- | --- | --- | --- | --- | --- | --- | --- |
| CD4+<br>(Cells/μl) | 14 Days | 120 | 110 | 120 | 120 | 100 | 120 |
|  | 28 Days | 470 | 450 | 370 | 320 | 230 | 260 |
|  | 42 Days | 1200 | 1100 | 900 | 380 | 540 | 700 |
|  | 56 Days | 900 | 830 | 600 | 385 | 500 | 560 |
|  | Normal Range | 359-1954 |  |  |  |  |  |
| CD8+<br>(Cells/μl) | 14 Days | 900 | 830 | 770 | 120 | 820 | 880 |
|  | 28 Days | 660 | 600 | 420 | 120 | 700 | 780 |
|  | 42 Days | 340 | 390 | 420 | 80 | 630 | 570 |
|  | 56 Days | 380 | 320 | 400 | 78 | 600 | 460 |
|  | Normal Range | 220-1100 |  |  |  |  |  |
| CRP<br>(ng/ml) | 14 Days | 48 | 47 | 49 | 50 | 44 | 45 |
|  | 28 Days | 5.8 | 6.5 | 4.6 | 66 | 29 | 22 |
|  | 42 Days | 89 | 92 | 22 | 100 | 122 | 105 |
|  | 56 Days | 4 | 11 | 22 | 140 | 0.9 | 2 |
|  | Normal Range | 0.78-25 |  |  |  |  |  |
| D-dimer<br>(ng/ml) | 14 Days | 66 | 64 | 65 | 69 | 65 | 64 |
|  | 28 Days | 36 | 42 | 47 | 42 | 55 | 50 |
|  | 42 Days | 28 | 33 | 37 | 59 | 55 | 49 |
|  | 56 Days | 22 | 26 | 33 | 67 | 20 | 15 |
|  | Normal Range | 0.6-50 |  |  |  |  |  |
| IL-6<br>(ng/ml) | 14 Days | 38 | 34 | 36 | 1.3 | 40 | 38 |
|  | 28 Days | 0.3 | 0.4 | 0.7 | 2.1 | 1.4 | 1 |
|  | 42 Days | 0.18 | 0.23 | 0.32 | 2.9 | 0.34 | 0.24 |
|  | 56 Days | 0.1 | 0.14 | 0.02 | 23 | 0.44 | 0.19 |
|  | Normal Range | 0.01-0.2 |  |  |  |  |  |
| Qualitative<br>IgM | 14 Days | Positive | Positive | Positive | Positive | Positive | Positive |
|  | 28 Days | Positive | Positive | Positive | Positive | Positive | Positive |
|  | 42 Days | Positive | Positive | Positive | Positive | Positive | Positive |
|  | 56 Days | Negative | Negative | Positive | Negative | Negative | Positive |
| Qualitative<br>IgG | 14 Days | Negative | Negative | Negative | Negative | Negative | Negative |
|  | 28 Days | Negative | Negative | Negative | Negative | Negative | Positive |
|  | 42 Days | Positive | Negative | Negative | Negative | Positive | Positive |
|  | 56 Days | Positive | Positive | Negative | Positive | Positive | Positive |
| Qualitative<br>RNA | 14 Days | Negative | Negative | Negative | Negative | Negative | Positive |
|  | 28 Days | Negative | Negative | Negative | Negative | Negative | Positive |
|  | 42 Days | Negative | Negative | Negative | Positive | Negative | Negative |
|  | 56 Days | Negative | Negative | Negative | Positive | Negative | Negative |

**Figure 16:**
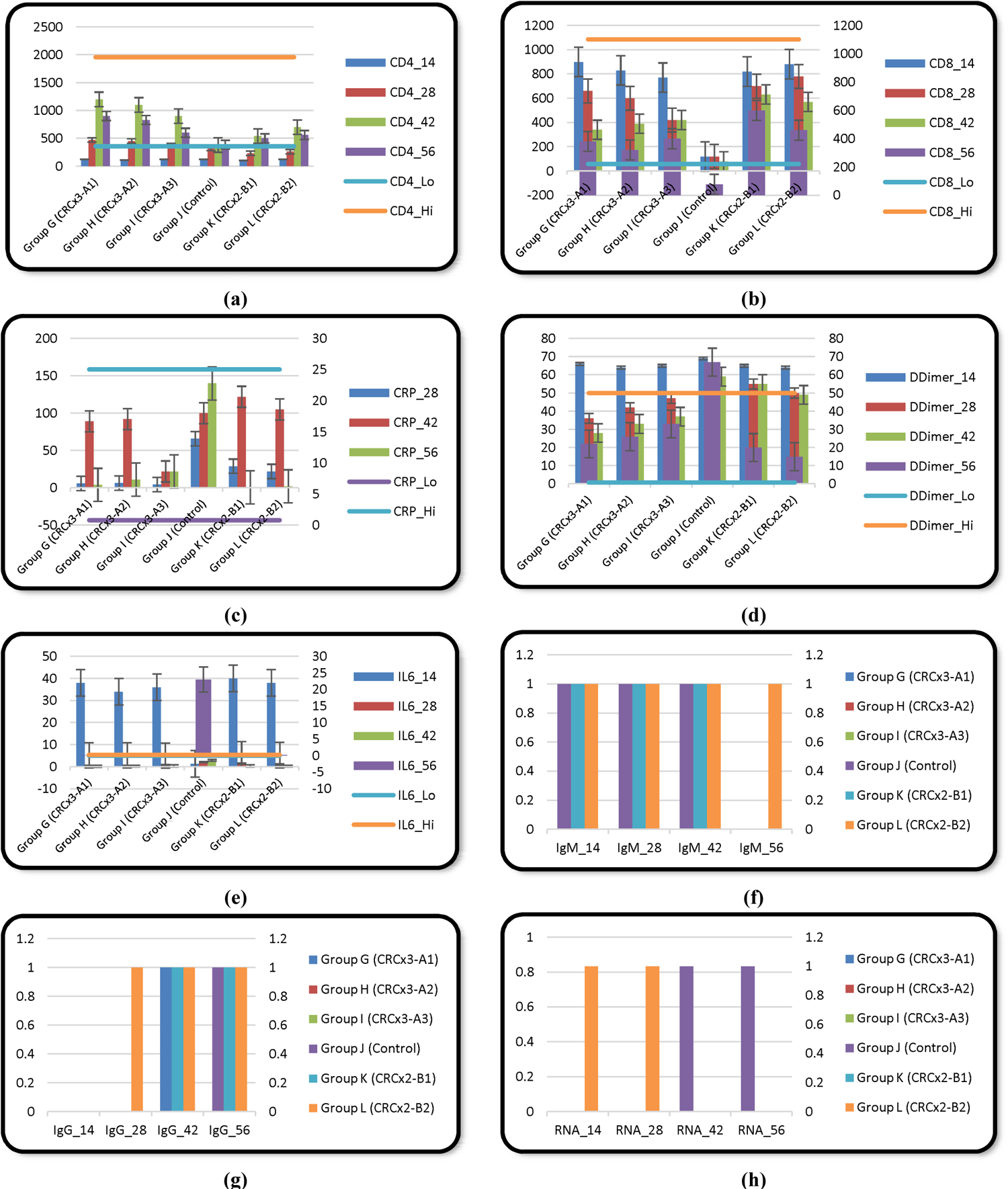
Quantitative and Qualitative Immunological and Biochemical Assays for Therapeutic Approach Groups on Days 14, 28, 42 & 56 with CRCx3 (Groups G, H & I), CRCx2 (Groups (K & L), and Control (J): (a) Quantitative CD4+. (b) Quantitative CD8+. (c) Quantitative C-Reactive Proteins (CRP). (d) Quantitative D-dimer. (e) Quantitative IL-6. (f) Qualitative SARS-Cov-2 IgM. (g) Qualitative SARS-Cov-2 IgG. (h) Qualitative SARS-Cov-2 RNA. Note: In quantitative assays, the horizontal lines represent the low and high normal values. In qualitative assays, 1=Positive; 0=Negative.

**Table 8:** Table 3: Neutralizing antibodies (NAbs) titers on days 14, 28, 42, and 56 in therapeutic regimens.

| Animal Groups | Days | Group G | Group H | Group I | Group J | Group K | Group L |
| --- | --- | --- | --- | --- | --- | --- | --- |
|  | Regimen | CoV-2 +<br>CRCx3<br>(0.4x3) | CoV-2 +<br>CRCx3<br>(0.35x3) | CoV-2 +<br>CRCx3<br>(0.25x3) | CoV-2 +<br>Nil | CoV-2 +<br>CRCx2<br>(0.4x2) | CoV-2 +<br>CRCx2<br>(0.25x2) |
| Anti-A1<br>(ng/ml) | 14 Days | 0 | 0 | 0 | 0 |  |  |
|  | 28 Days | 400 | 340 | 300 | 0 |  |  |
|  | 42 Days | 950 | 800 | 710 | 0 |  |  |
|  | 56 Days | 600 | 520 | 480 | 0 |  |  |
|  | Normal Range |  |  |  | 0.2-1 |  |  |
| Anti-A2<br>(ng/ml) | 14 Days | 0 | 0 | 0 | 0 |  |  |
|  | 28 Days | 230 | 200 | 140 | 0 |  |  |
|  | 42 Days | 450 | 410 | 350 | 0 |  |  |
|  | 56 Days | 320 | 270 | 200 | 0 |  |  |
|  | Normal Range |  |  |  | 0.5-1 |  |  |
| Anti-A3<br>(ng/ml) | 14 Days | 0 | 0 | 0 | 0 |  |  |
|  | 28 Days | 450 | 380 | 300 | 0 |  |  |
|  | 42 Days | 750 | 640 | 575 | 0 |  |  |
|  | 56 Days | 500 | 400 | 310 | 0 |  |  |
|  | Normal Range |  |  |  | 0.5-5 |  |  |
| Anti-B1<br>(ng/ml) | 14 Days |  |  |  | 0 | 100 | 85 |
|  | 28 Days |  |  |  | 100 | 1200 | 1000 |
|  | 42 Days |  |  |  | 200 | 950 | 870 |
|  | 56 Days |  |  |  | 360 | 600 | 520 |
|  | Normal Range |  |  |  | 0.5-2 |  |  |
| Anti-B2<br>(ng/ml) | 14 Days |  |  |  | 22 | 30 | 24 |
|  | 28 Days |  |  |  | 120 | 600 | 520 |
|  | 42 Days |  |  |  | 180 | 500 | 450 |
|  | 56 Days |  |  |  | 300 | 430 | 400 |
|  | Normal Range |  |  |  | 0.2-2 |  |  |

**Figure 17:**
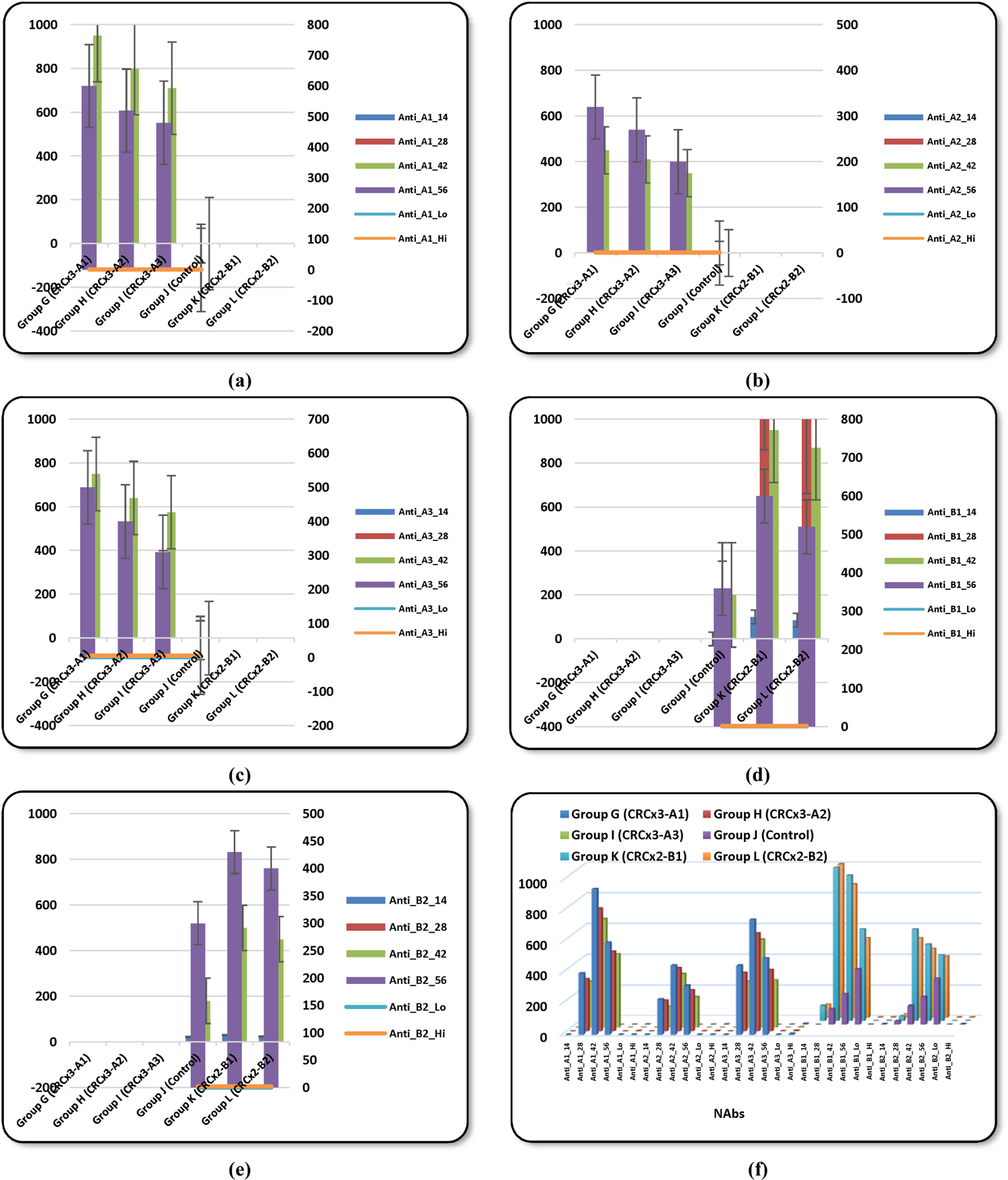
Different neutralizing antibodies (NAbs) in Therapeutic Approach Groups on Days 14, 28, 42 & 56 with CRCx3 (Groups A, B & C), CRCx2 (Groups (E & F) and Control (D): (a) Anti-A1. (b) Anti-A2. (c) Anti-A3. (d) Anti-B1. (e) Anti-B2. (f) All NAbs. Note: The horizontal lines represent the low and high normal values.

